# TOC1 phosphorylation disproportionally enhances chromatin binding at rhythmic gene promoters

**DOI:** 10.1101/2025.01.09.632189

**Authors:** Jiapei Yan, Ying Zhang, Guoting Chen, Cuilin Gui, Zhangli Tu, Donglin Cao, Xingwang Li, David E Somers

## Abstract

Protein phosphorylation is a key regulatory mechanism in circadian systems. TIMING OF CAB EXPRESSION 1(TOC1) is a core transcriptional repressor in the plant circadian system that is phosphorylated near its N-terminus. Phenotype testing of single and multi-phosphosite mutants shows incomplete rescue of the short period *toc1* mutant. We establish that TOC1 phosphorylation (particularly at S175) is necessary for optimal interaction with FAR-RED ELONGATED HYPOCOTYL3 (FHY3) and PHYTOCHROME INTERACTING FACTOR 5 (PIF5) at the *CIRCADIAN CLOCK-ASSOCIATED 1 (CCA1)* promoter to downregulate *CCA1* expression. At the same time, expression of the closely related *LATE ELONGATED HYPOCOTYL (LHY)* also requires TOC1, but is independent of the TOC1 phosphorylation state, suggesting different TOC1-dependent mechanisms in the repression of these two genes. We additionally show how phosphorylation-dependent interactions of TOC1 at specific clock gene promoters selectively regulate these circadian system components more acutely than non-rhythmic genes. Our genome-wide analysis shows that the phosphostate of TOC1 is important for optimal chromatin presence and robust rhythmic gene expression.

## Introduction

Adapting to the day and night cycles that arise from the rotation of the earth, most organisms have developed an endogenous oscillator, the circadian clock, to synchronize their biological processes with a rhythm of ca. 24 h (*1–3*). Plants as sessile organisms, particularly rely on this internal clock to anticipate the external cyclic changes and adjust their growth, physiology and development in advance (*4, 5*).

The molecular mechanism of circadian clock in plants was described as an autoregulatory transcriptional feedback network (*6*). The central oscillator of *Arabidopsis* clock is composed of two core groups of clock components. One group contains two morning-expressed MYB-like transcription factors, CIRCADIAN CLOCK-ASSOCIATED 1 (CCA1) and LATE ELONGATED HYPOCOTYL (LHY). These two closely-related morning-phased transcription factors bind the evening element (EE) of evening-phased clock genes, such as *TIMING OF CAB EXPRESSION 1* (*TOC1*), *LUX ARRHYTHMO (LUX)*, *EARLY FLOWERING 3 (ELF3)* and *ELF4* and repress their transcription (*7–12*). PSEUDO-RESPONSE REGULATOR (PRR) proteins, including TOC1, PRR3, PRR5, PRR7 and PRR9 comprise another core group of clock components in *Arabidopsis*, which sequentially repress *CCA1* and *LHY* expression and thus form transcriptional feedback loops (*13, 14*) that sustain the pace of the central oscillator.

A key feature of circadian systems is synchronization with the surrounding environment through informational input pathways. Light is one of the major signals shaping the rhythmic expression of core clock genes and thus resets the clock (*15, 16*). FAR-RED ELONGATED HYPOCOTYL3 (FHY3) acts as phytochrome signaling integrator and plays an important role in gating red light to the clock system in the morning (*17*). FHY3 and its paralog FAR-RED IMPAIRED RESPONSE1 (FAR1) are required for the light induction and normal rhythmic expression of *CCA1* through direct promoter binding (*18*). In contrast, PHYTOCHROME INTERACTING FACTOR5 (PIF5) and TOC1 antagonize FHY3 activity through physical interactions and repress *CCA1* expression from pre-dusk to midnight (*18*). Blue light signaling also plays a role in *CCA1* regulation. Cryptochrome 2 (CRY2) was found to form photobodies with TEOSINTE BRANCHED1-CYCLOIDEA-PCF 22 (TCP22) and activates *CCA1* expression through LIGHT REGULATED WDs (LWDs) and the TBS motif (*19*) in a blue light-dependent manner. Additionally, PHOTOREGULATORY PROTEIN KINASES 1 (PPK1) phosphorylates TCP22 and enhances the formation of CRY2-TCP22 photobodies to promote the expression of *CCA1* (*19*). These findings indicate a close relationship between light signaling components and clock proteins in coordinating robust rhythmic expression of *CCA1*.

Numerous studies in mammals, *Drosophila*, and *Neurospora* have shown the fundamental importance of post-translational regulation, particularly phosphorylation, in the circadian clock (*20, 21*). In Arabidopsis, phosphorylation also comprises an essential integral part of the circadian regulatory system (*22, 23*). For example, all PRR proteins are phosphorylated, and phosphorylation enhances their heterodimerization and possibly interactions with other proteins as well (*24*). A chemical screen of altered circadian period identified CKL4, a member of CASEIN KINASE 1 LIKE (CKL) family, that is possibly involved in phosphorylation of TOC1 and PRR5 (*25*). TOC1 and PRR5 are the most closely-related members of the PRR family, and PRR5 promotes TOC1 phosphorylation and nuclear transport through heterodimerization with the TOC1 N-terminus (*26*). Additionally, specific TOC1 N-terminal phosphosites are essential for NF-Y-TOC1 complex formation and its regulated photoperiodic hypocotyl elongation (*27*). However, how TOC1 phosphorylation regulates its role in the central oscillator is still unknown.

In this study we establish that N-terminal phosphorylation of TOC1, particularly serine 175 (S175), is required for sustaining both wild-type circadian period and the robustness of circadian oscillations. FHY3 interacts strongly with phosphorylated TOC1 and selectively shapes *CCA1* expression rhythm, but not *LHY*, through the *FBS* element. Loss of either S175 phosphorylation or FHY3 results in shortened period and decreased amplitude, while absence of both leads to arrhythmicity.

Genome-wide transcriptome analyses identified global changes in amplitude and phase of rhythmically expressed genes between *TOC1* and *5X* lines. Our findings suggest that these changes in circadian oscillations of gene expression are likely due to the diminished 5X binding to central oscillator clock genes. Additionally, we identify a very different categories of non-cycling TOC1 target genes that point to possible non-circadian roles for TOC1. Together, our results expand our understanding of post-translational regulation of the circadian clock and provide a comprehensive analysis of how TOC1 phosphorylation affects genome rhythmicity and interesting insights into the selective regulation of *CCA1* by TOC1 interacting with different co-factors.

## MATERIALS AND METHODS

### Plant Materials and Growth Conditions

The wild-type and all mutants of *Arabidopsis thaliana* plants used in this study were of the Colombia-0 (Col-0) ecotype. The *toc1-101* (*toc1*, (*28*)), *pif3-3 pif4-2 pif5-3 toc1-101* (*pif345toc1*,), , *prr5-1 prr7-1* (*prr5prr7*, (*29, 30*)), *nf-yc3-1 nf-yc4-1 nf-yc9-1* (*nf-yc349*, (*31*)) and *nf-yc3-1 nf-yc4-1 nf-yc9-1 toc1-101* (*nf-yc349toc1*,(*27*)) mutant lines, as well as *CCA1:LUC* line (Salome & McClung, 2005) were previously described. The *fhy3-11* (SALK_002711) (*18*) was obtained from the Arabidopsis Biological Resource Center (http://www.arabidopsis.org/). The *CCA1:LUC* line and TOC1 native promoter lines *fhy3-11* to produce *fhy3 toc1* double mutant, and the *CCA1:LUC* line was crossed with various mutants (*27*) was crossed into *fhy3*and *pif3/4/5*, respectively. The point mutations in 5X (T135A/S175A/S194A/S201A/S204A), 135A, and 175A lines were introduced by site-directed mutagenesis driven by the *TOC1* native promoter as described previously (*27*). Homozygous F3 progenies of each plant line were used for bioluminescence and ChIP assays.

Arabidopsis seeds were surface sterilized and stratified at 4°C for 4 days. Arabidopsis seedlings used in this study were all grown at 22°C. For bioluminescence acquirement and period estimate, seedlings were entrained under 12h white fluorescent light /12h dark cycles for 7 days on MS (Murashige and Skoog) plates with 3% sucrose and 0.8% agar as previously described (*32*). Seedlings were then subjected to constant white (30 µmol m^−2^ s^−1^) light treatment conducted in a Percival E30LEDL3 growth chamber (Percival Scientific, Perry, IA). For protein immunoblotting, transcript abundance analyses and ChIP assays in 12h white fluorescent light (50 µmol m^−2^ s^−1^)/12h dark cycles, for 10 days on MS plates with 3% sucrose and 0.8% agar. For transient expression experiments in *N. benthamiana*, 4-week-old plants were infiltrated with *Agrobacteria* as described earlier (*24*). Tissues were harvested on the third day at ZT12 (16 h light/8 h dark).

### Constructs

To construct TAP-tagged TCPs, FHY3, FAR1 and PIF5, full-length coding region was amplified and each amplicon was subcloned into pENTR/D-TOPO and placed upstream of the TAP tag by LR recombination (Invitrogen) into the binary vector pYL436 (ABRC; CD3–679). To generate TOC1-GFP, 5X-GFP, T135A-GFP, S175A-GFP and S175D-GFP for Co-IP assays in *N. benthamiana*, the cDNA fragments of each gene were subcloned into pENTR2B or pENTR/D-TOPO and placed into the binary vector pCsVMV-GFP-N-1300 by LR recombination (*26*). To generate plasmids for transient expression in protoplasts, the cDNA fragments of TOC1 and 5X were transferred from pENTR/D-TOPO entry clones to pCsVMV:GFP-C-999 destination vector by LR recombination.

### Protein Extraction and Immunoblots

Total protein extraction was carried out as described earlier (*24, 33*). For the immunoblot analyses, proteins were separated by 8-10% SDS–PAGE (acrylamide:bisacrylamide 37.5:1) for regular protein electrophoresis or 8% SDS– PAGE (acrylamide:bisacrylamide 150:1) to differentiate wild type and phosphosite mutants of TOC1. Immunoblotting was performed using 1:5000 dilution of primary polyclonal anti-GFP (Abcam ab6556), 1:10000 dilution of polyclonal ADK primary antibodies (gift from Dr David Bisaro) followed by ECL detection using anti-rabbit IgG with horseradish peroxidase (HRP) linked whole antibody (GE healthcare NA934V). For immunoblot of primary anti-Myc (Sigma-Aldrich M4439, 1:4000 dilution), HRP-linked anti-mouse IgG (Sigma-Aldrich, A0198) was used as secondary antibody. Chemiluminance reactions were performed with Supersignal West Pico and Femto Chemiluminescent Substrates (Thermo Scientific).

### Co-immunoprecipitation (Co-IP) Assays

Co-IP assays were conducted as previously described (*27*). Briefly, agrobacteria containing corresponding expression clones were coinfiltrated into 4-wk-old *N. benthamiana* leaves, the total proteins were extracted and used for immunopreciptation with Human IgG-Agarose (Sigma-Aldrich, A6284). The immune complex was washed and subsequently released from resin by HRV-3C protease digestion. The eluted TAP-and GFP-proteins were detected by Western blotting using Myc antibody (Sigma-Aldrich, M4439) and GFP antibody (Abcam, ab6556), respectively.

### Chromatin Immunoprecipitation (ChIP) Assays

ChIP assays were conducted as previously described. Briefly, 10-day-old Arabidopsis seedlings grown in L/D conditions were harvested and cross-linked with 1% formaldehyde by vacuum infiltration. Cross-link was then quenched by 0.125 M glycine. ChIP extracts were incubated with anti-GFP (Abcam ab6556) overnight and immunoprecipitated by Protein G Dynabeads (Thermo scientific 10004D). DNA of input and reverse cross-linked IP samples was purified using QIAquick PCR clean up Kit (Qiagen) and subjected to quantitative PCR using promoter specific primers (*34*). To perform ChIP assays using protoplasts, pCsVMV:GFP-C-999 (express GFP alone) was used for plasmid construction and the ChIP assays was performed as previously described (*27*).

### Luminescence Measurement and Circadian Rhythm Analysis

Luminescence acquirement was conducted as previously described (*32*). Data analysis and period estimate (based on *CCA1:LUC* reporter) were performed as in (*35*).

### RNA Extraction and Quantitative PCR

10-day-old seedlings grown in L/D conditions were harvested at indicated time points. Total RNA extraction, DNase treatment, cDNA synthesis and quantitative PCR were done as described earlier (*35*).

### ChIP-seq libraries preparation

10-day-old Arabidopsis plants were harvested at ZT14, crosslinked with 1% formaldehyde and quenched with 0.2 M glycine. For each experiment, about 0.5 to 1g of the sample was ground into fine powder in liquid nitrogen. Nuclei were lysed in buffer S (50 mM HEPES-KOH, 150 mM NaCl,1 mM EDTA, 1% Triton X-100, 0.1% sodium deoxycholate, 1% SDS, 2 mM NaF, 2mM Na_3_VO_4_, 50 µM MG132, 50 µM MG115, 50 µM ALLN and protease inhibitor cocktail) at 4 °C, and chromatin was fragmented into 200-600 bp by ultrasound processing using a Bioruptor (Diagenode). The lysates were centrifuged at 20,000 g at 4 °C for 10 min. Then 5-10 μL GFP antibody (Abcam, ab6556) and 200 μL suspended protein G magnetic beads (Life Technologies, 10003D) were incubated at 4 °C for 1 to 6 h and rotated to generate antibody-bead complexes. The protein-DNA complexes were subjected to immunoprecipitation by incubating the antibody-bead complexes with the fragmented chromatin. Then, the target protein-DNA complexes were eluted from the beads by adding 100 μL of freshly prepared ChIP elution buffer (50 mM Tris-HCl pH 7.5, 10 mM EDTA, and 1% SDS). After reverse cross-linking of DNA and protein, ChIP-DNA was extracted using phenol: chloroform: isoamyl alcohol (P3803; Sigma-Aldrich), precipitated with ethanol, and resuspended in TE buffer. ChIP-DNA libraries were prepared using a NEBNext® Ultra™ II DNA Library Preparation Kit. AMPure XP beads (Beckman, A63881) were used to select 250-650 bp library fragments. Finally, the DNA fragments were sequenced using an Illumina HiSeq X Ten system (paired-end 150-bp reads).

### Whole transcriptome sequencing (RNA-seq) libraries preparation

RNA was isolated from 10-day-old arabidopsis plants grown under L/D conditions using RNeasy Plant Mini Kit (QIAGEN, 74904) according to the manufacturer’s instructions. 1.5 µg of total RNA was depleted of ribosomal RNAs using TruSeq Stranded Total RNA with Ribo-Zero Plant (Illumina, RS-122-2401) for RNA-seq according to the manufacturer’s instructions. RNA sequencing was performed on an Illumina HiSeq X Ten system (paired-end 150 bp reads).

### ChIP-seq data analysis

Raw data was checked for quality and filtered by FastQC and Trimmomatic, respectively. Clean reads were then mapped to the tair10 reference genome with BWA-MEM (version 0.7.17), using default parameters (*36*). After removing potential PCR repeats with Samtools (version 1.9) ,(*37*) peak calling was performed with MACS (version 2.1.1) (-f BAM -g 1.2e+8) (*38*). If RSC>1, NSC>1 and FRIP>40% of the obtained peak, it can be used for subsequent analysis. ChIP-seq peaks were visualized by Integrative Genomics Viewer (IGV) (version 2.5.0)(*39*) and the input file is in bigWig format, which can be converted by bamcoverage in deeptools (version 3.2.1) (*40, 41*).

### RNA-seq data analysis

Raw data of RNA-seq were first evaluated by FASTQC, and qualified data were filtered out by Trimmomatic (version0.36) (*42*). The filtered reads were mapped to the tair10 reference genome using Hisat2 (version 2.1.0)(*43*), which requires the use of the RNA-strandness RF parameter as the data was strand-specific library building. Gene expression was quantified using HTseq. If the sum of the expression of the same gene in all samples is greater than 5, the gene is defined as expressed and used for subsequent analyses (*41*).

### Rhythmicity characterization using time-series RNA-seq data

Rhythmicity prediction was performed as previously described (*44*). Briefly, RNA-seq data was normalized at all time points, BIO_CYCLE was used to predict rhythmicity of the expressed genes. The cycle parameter is set between 20 and 28. The rhythmicity of the gene was determined according to p_value and relative entropy error. If the p_value is less than 0.05 and the relative entropy error is closer to 1, the gene was considered as rhythmic. The differentially oscillated genes between *TOC1* and *5X* plants were determined by phase change > 2 h, amplitude change > 1.5-fold.

The sinusoidal model fitting for the RNA-seq traces was performed by using the ggplot2 package in R on the six-time-point expression data of RNA-seq with three replicates (*45*). The fitting formula for the sinusoidal model is cos(2*pi*Time/24) + sin(2*pi*Time/24). Since the fitted sinusoidal function is cyclical, providing expression data at six time points allows for the plotting of diurnal expression curves for each gene. The data point in the figure shows the average of the three replicates, and the confidence interval of the curve was determined by the geom_smooth parameter.

### KEGG pathway analysis

The plant GeneSet Enrichment Analysis Toolkit was used in KEGG pathway enrichment analysis. The FDR<0.05 was set as the cutoff for statistical significance. http://structuralbiology.cau.edu.cn/PlantGSEA/analysis.php (*46*).

### Motif analysis

Motif enrichment analysis was done by Homer software as previously described (*37, 47*). The parameter “-mset plants” was added for plant motif analysis and other parameters are default.

## RESULTS

### TOC1^S175^ is essential for its function in the central oscillator

Our previous study identified 5 phospho-sites at N-terminus of TOC1, mutations in all of which (alanine substitutions; 5X) result in poor interactions with NF-Y co-factors and increased hypocotyl growth under short days (*27*). In addition to the growth phenotype, TOC1 5X also showed significantly shortened circadian period compared to wild-type (*27*). To investigate the role of TOC1 phosphorylation and NF-Ys in the context of the circadian oscillator, we first examined the clock-related phenotypes in various TOC1 phospho-site substitutions in either *toc1* single or *nf-yc3/4/9 toc1* quadruple mutant backgrounds under constant white light. We found 5X and TOC1^S175A^ both exhibit a 1.5h shorter period (Fig.1A and Supplemental Fig.1), increased relative amplitude error (RAE) (Fig.1C) and *CCA1-LUC* oscillation that dampens rapidly relative to WT (Fig.1B and E), indicating phosphorylation at TOC1^S175^ is essential for normal circadian oscillations, as we previously reported (*27*). In contrast, mutation of the closely positioned T135 residue to alanine (TOC1^T135A^)(*27*) had no effect on period. Consistent with this result, the protein oscillations of only 5X and TOC1^S175A^ are phase advanced under 12 h light/ 12 h dark cycles (Supplemental Fig. 2).

**Figure 1.**
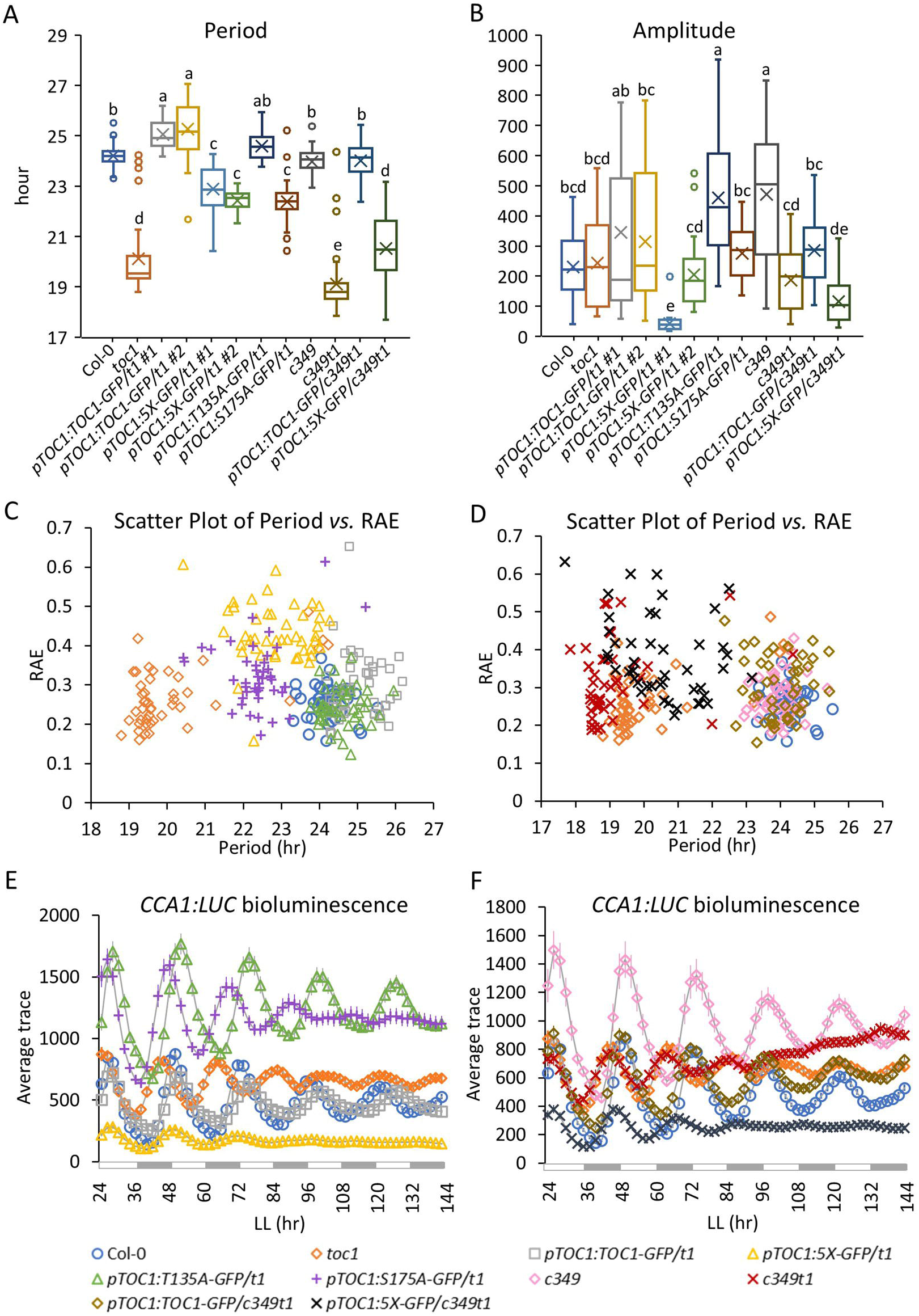
Shorter period and dampened *CCAJ-LUC* oscillations in *5X* and *S175A* mutants. Free-running period **(A),** amplitude **(B),** RAE **(C, D)** and average *CCAJ-LUC* bioluminescence **(E, F)** of Col-0, *tocl, nfyc3/4/9 (c349), nfyc3/4/9 tocl (c349tl)* and *TOCJ* native promoter lines in *tocl (tl)* single or *nf-yc3/4/9 tocl* quadruple mutant backgrounds. #1 and #2 indicate the two independent lines of the corresponding transgene. In the boxplots **(A, B),** the middle line of the box represents the median, x represents the mean, the bottom line and the top line indicate the 1st and 3rd quartile, respectively. The whiskers extend from the ends of the box to the minimum value and maximum value. In the scatterplot **(C, D),** data points from Col-0 and the four *TOCJ* native promoter lines in *tocl* mutant background are circled by dashed lines. Seedlings were entrained in 12-h/12-h light/dark cycles for 7 days and then transferred to constant white light at ZT2 for image acquisition at 2-h intervals for 1 week. 45 seedlings from 3 independent trials were averaged (n=45). Different letters indicate statistically significant differences *(p* < 0.01, one-way ANOVA followed by Tukey-Kramer HSD test).

The period of the *nf-yc3/4/9* triple mutant is not different from Col-0, and the period of the *nf-yc3/4/9 toc1* quadruple mutant (Supplemental Table 1) is similar to *toc1* (Fig.1A, D and Supplemental Fig. 1C), supporting the notion of a TOC1-anchored trimeric complex at the chromatin, comprised of NF-YB/C factors as previously reported for hypocotyl length control (*27*). 5X in *nf-yc3/4/9 toc1* was unable to restore clock-related phenotypes and showed a short period (Supplemental Table 1) (Fig.1A), high RAE (Fig.1D) and strongly dampened *CCA1-LUC* oscillations, similar to the 5X *nf-yc3/4/9* triple mutant (Fig.1F).

### Phosphorylation stabilizes TOC1 chromatin residence at clock genes

To investigate how TOC1 phosphorylation affects the control of central oscillator genes, we assayed the residence of both TOC1 and 5X at the promoters of several key clock genes previously reported as binding TOC1 (*48*). We chose ZT14 for ChIP assays since TOC1 showed equal protein levels in both *pTOC1:TOC1-GFP* and *pTOC1:5X-GFP* at this time point under 12h/12h light/dark cycles (Supplemental Fig. 3A).

In most cases the 5X mutations diminished TOC1 presence at the genes tested. *PRR7*, *PRR9* and *GI* showed 30-40% less TOC1 at their promoters and an almost complete loss of TOC1 at *CCA1* in *pTOC1:5X-GFP* line was observed (Fig.2A). However, there were no differences between TOC1 and 5X binding at the *LHY* promoter (Fig.2A). These results suggest that phosphorylation enhances TOC1 chromatin binding activity at a subset of target clock genes, such as *CCA1*, *PRR7*, *PRR9* and *GI*.

**Figure 2.**
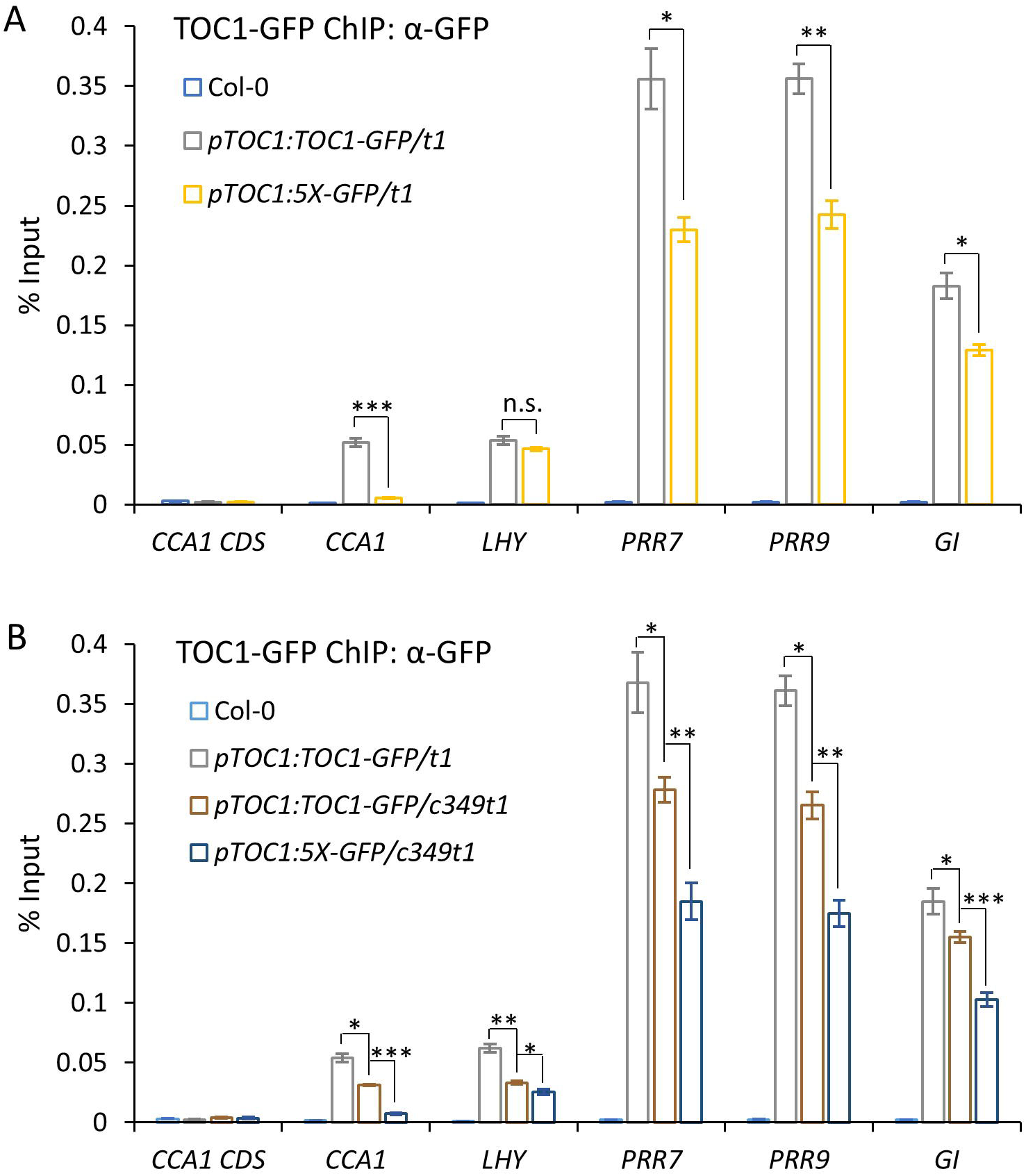
Robust TOCl chromatin residence depends on TOCl phosphostate and NF-YC proteins. ChIP-qPCR of TOC1 binding to the promoter region of several core clock genes in Col-0, wild-type *TOCJ* and *5X* in *tocl (ti)* single **(A)** or *nf-yc3/4/9 tocl (c349tl)* quadruple **(B)** mutant backgrounds. 10-d-old seedlings grown at 12-h/12-h light/dark conditions were harvested at ZT14 **(A)** or ZT13 **(B),** TOCl-GFP was immunoprecipitated by a-GFP with magnetic protein G beads. Data were averaged from 3 independent trials, each trial with 2 technical repeats. Error bars indicate SEM. Asterisks indicate significant differences *(*p* < 0.05, *\*\*p* < 0.001, *\*\*\*p* < 0.0001, one­ way ANOVA followed by two-tail Student’s t-test), n.s. = not significant.

We next examined whether *NF-YC*s play a role in TOC1 presence at target clock genes. These experiments were conducted at ZT13 to ensure similar TOC1 levels in the respective genotypes (Supplemental Fig. 3). ChIP of TOC1 in an *nf-yc3/4/9* background showed a 20-40% reduction in chromatin occupancy, relative to WT, at all tested clock promoters. The phosphorylation defects in 5X decreased chromatin binding by a further, except for *LHY* (Fig. 2B). Similar to in the *toc1* background (Fig. 2A), chromatin residence of TOC1 and 5X in the *nf-yc3/4/9 toc1* quadruple mutant at the *LHY* promoter were very similar, whereas 5X binding to *CCA1* promoter diminished to almost background levels (Fig. 2B). These results suggest that NF-YCs are not the factors responsible for the striking difference between TOC15X and TOC1WT binding at the *CCA1* and *LHY* promoters. Together with the marginal period phenotype of *nf-yc3/4/9*, these results exclude the three *NF-YC* genes as major determinants in regulating phosphorylation-dependent functions of TOC1 in the circadian system.

### Phosphorylation of TOC1^S175^ promotes *CCA1* oscillation

To test if TOC1^S175^ is the most essential TOC1 phospho-site, we next performed ChIP with *pTOC1:T135A-GFP* and *pTOC1:S175A-GFP*. Similar to *5X*, *S175A* showed strikingly reduced residence at *CCA1*, whereas *T135A* binding was only marginally less at *CCA1* relative to wild type (Fig 3A). The differences in TOC1 chromatin residence were not due to differences in protein abundance between plant lines (Supplemental Fig. 2). In contrast to *CCA1*, all 4 different TOC1 phospho-variants showed similar presence at the *LHY* promoter (Fig.3A).

**Figure 3.**
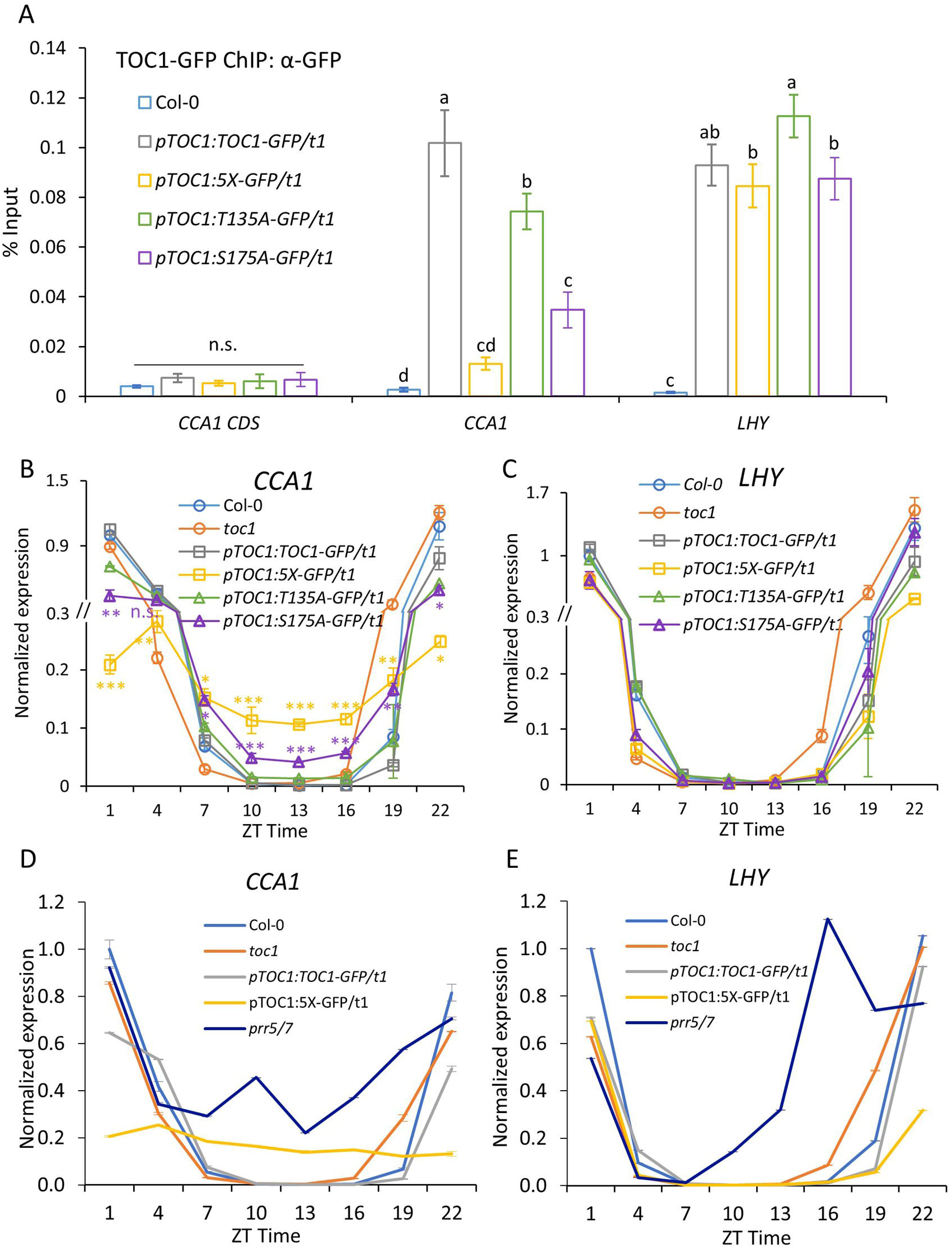
TOCl phosphomutants show reduced binding to *CCAJ* and diminished *CCAJ* oscillations. **A.** ChIP-qPCR ofTOCl binding to the promoter region of *CCAJ* (- 730 to -648) and *LHY* (-1062 to -958) at ZT14 in Col-0, wild type and phospho-mutant lines of *TOCJ.* Different letters indicate significant differences (p < 0.05, one-way ANOVA followed by two-tail Student’s t-test), n.s. = not significant. **B, C.** Time-series expression of *CCAJ* **(B)** and *LHY* (C) in Col-0, *tocl,* wild type and phospho-mutant lines of *TOCJ.* 10-d-old seedlings grown at 12-h/12-h light/dark conditions were harvested at indicated time points. Data were averaged from 3 independent trials, each trial with 2 technical repeats. Error bars indicate SEM. Asterisks indicate significant differences of 5X and Sl 75A compared to wild-type TOCl at each time point *(*p* < 0.05, *\*\*p* < 0.001, *\*\*\*p* < 0.0001, Student’s t-test), n.s. = not significant. **D, E.** Time­ series expression of *CCAJ* **(D)** and *LHY* **(E)** in Col-0, *prr5 prr7 (prr5/7)* and indicated plant lines. Error bars indicate SEM of two repeats. 10-d-old seedlings grown at 12-h/12-h light/dark conditions were harvested at indicated time points. Expression was normalized to Col-0 at ZTl.

To determine the consequences of reduced 5X binding on gene expression, we next examined transcript levels of *CCA1* and *LHY* in Col-0, *toc1* and the 4 *TOC1* phospho-variant lines over a 12/12 light/dark cycle. Interestingly, *CCA1* displayed significantly lower amplitude oscillations in *5X* and *TOC1^S175A^* plants than in Col-0, *toc1* and *TOC1^T135A^* lines (Fig. 3B). These results indicate that these phosphorylation defective mutants do not simply phenocopy the *toc1* null mutant, but rather they may be neomorphic in the context of *CCA1* transcription. We hypothesize that 5X and TOC1^S175A^ may act as dominant negative mutants by interfering with the binding of other PRR repressors to the *CCA1* promoter, possibly through heterodimerization. Consistent with this hypothesis, the *prr5prr7* double mutant (*prr5/7*) showed a strikingly higher trough of *CCA1* expression, particularly from ZT7 to ZT16 (Fig. 3D, E), similar to results reported previously (*14, 49*) . Sequestration of endogenous PRR5 and/or PRR7 by 5X or TOC1^S175A^ heterodimerization could account for the similar high trough phenotypes of the 5X and *prr5/7* mutants particularly between ZT7 and 10, where *CCA1* expression is high but *LHY* is not.

### Phosphorylation of TOC1^S175^ enhances binding affinity with FHY3 and PIF5 but not TCPs

To identify factors that work together with TOC1 to cause the differential transcriptional regulation between *CCA1* and *LHY*, we examine the promoter sequence of *CCA1* and *LHY* and identified two *cis*-elements that are present in *CCA1* but not in *LHY* : an FHY3/FAR1 binding sequence (FBS) and a TCP (TEOSINTE BRANCHED 1, CYCLOIDEA, PCF1) binding site (TBS) (Fig. 4A).

**Figure 4.**
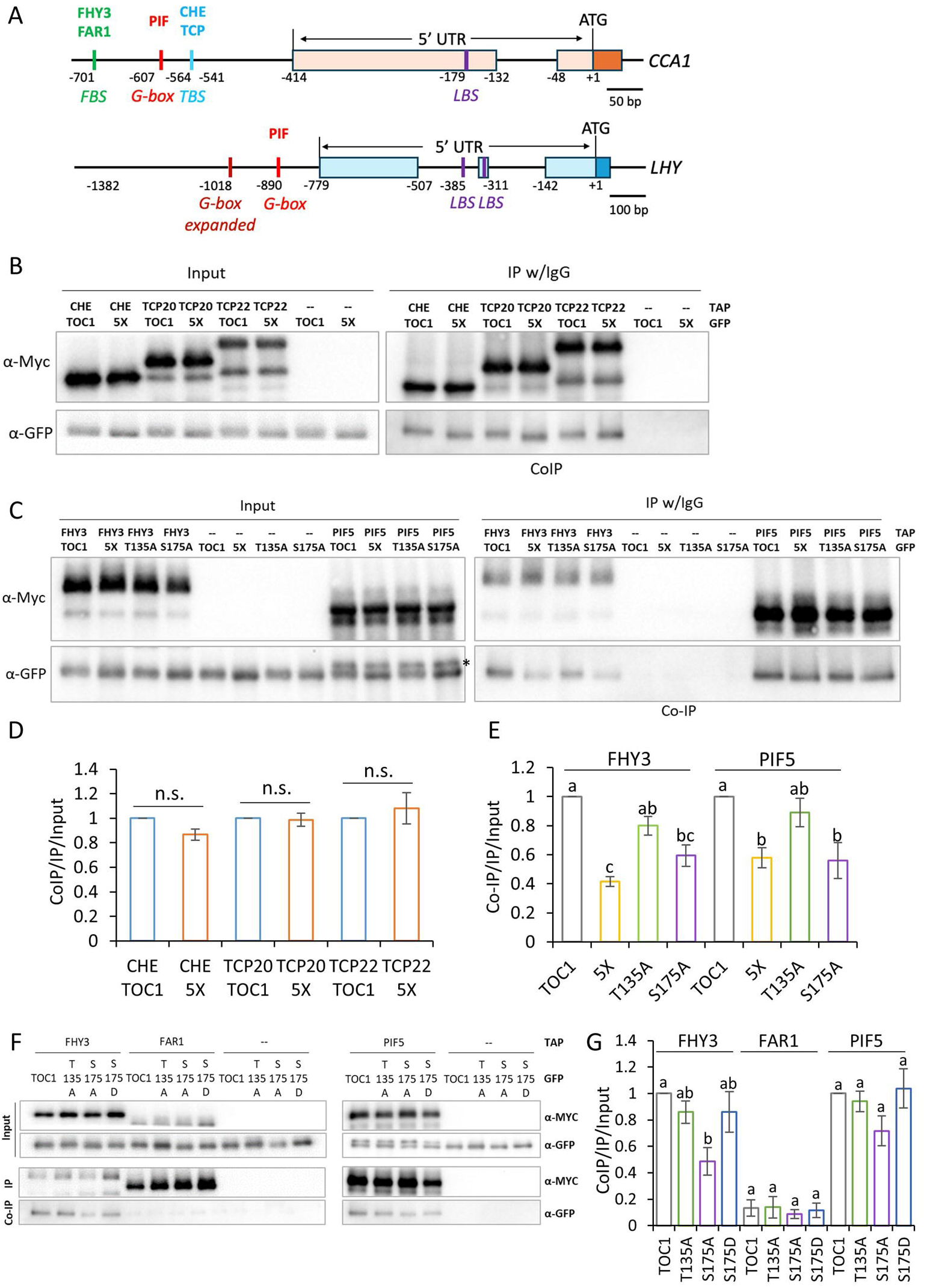
TOCl phosphomutants interact more poorly with FHY3 and PIFS. **A.** Schematic representation of the positions of various cis-elements in the CCAl and LHY promoter. FBS: FHY3/FAR1 binding sequence. TBS: TCP binding sequence. G-box: PIF binding motif. LBS: LUX binding sequence. **B, C.** Co-IP of TAP tagged CHE/TCP **(B),** FHY3 or PIF5 (C) with TOCl-GFP or TOCI carrying alanine substitutions in *N. benthamiana.* Protein extracts were immunoprecipitated by lgG resins followed by HRV-3C protease digestion. TAP tagged proteins were detected by anti-MYC and TOCl was detected by anti-GFP. Asterisks in C indicates PIF5 that was unspecifically recognized by the secondary a-Rabbit. **D.** Quantitation of protein interactions in Fig.4B, Supplemental Fig.4A and B. **E.** Quantitation of protein interactions in Fig.4C and Supplemental Fig.SA. For D and E, results from 3 independent experiments were averaged (except CHE with 2 trials). Error bars indicate SEM, different letters indicate significant difference *(p* < 0.05, one-way ANOVA followed by Tukey HSD test). **F.** Co­ IP of TAP tagged FHY3, FARl or PIF5 with TOCl-GFP and TOCl carrying alanine or aspartate substitution in *N. benthamiana.* Protein extracts were immunoprecipitated by IgG resins followed by HRV-3C protease digestion. TAP tagged proteins were detected by anti-MYC and TOCI was detected by anti-GFP. **G.** Quantitation of protein interactions in F and Supplemental Fig.SB. Results from 3 independent trials were averaged. Error bars indicate SEM, different letters indicate significant difference *(p* < 0.05, one-way ANOVA followed by Tukey HSD test).

Since phosphorylation affects TOC1 binding affinity with the NF-Y cofactors (*27*), we tested the interactions of TOC1 phospho-variants with FHY3/FAR1 and TCPs. CHE, TCP20 and TCP22 are members of the TCP family of transcription factors (*50*). CHE interacts with and recruits TOC1 to the *CCA1* promoter, while TCP20 and 22 form tetramers with LWD1/2 to activate *CCA1* expression (*51, 52*). However, none of the three TCP proteins show significant differences from WT in their interactions with TOC1 phospho-variants. (Fig. 4B, D and Supplemental Fig. 4).

We next tested FHY3 and PIF5 which interact with TOC1 and coordinately regulate *CCA1* expression under light/dark cycles (*18*). In these cases FHY3 and

PIF5 both showed a 60% and 40% reduced interaction with 5X and S175A, respectively, (Fig. 4C, E and Supplemental Fig. 5). In contrast, the phosphomimetic mutant S175D and wild-type TOC1 interacted similarly with FHY3 and PIF5 (Fig. 4F,G and Supplemental Fig. 5D), suggesting that the differential binding affinity between TOC1 phospho-variants and FHY3/PIF5 results from phosphorylation, not protein conformation. Our results are also consistent with a previous report mapping the TOC1-FHY3 interaction domain to a region that includes S175 (aa 141 to 520) (*18*). In contrast to FHY3, FAR1 interactions with TOC1 were negligible (Fig 4F,G; Supplemental Fig. 5).

Despite the diminished binding of 5X and S175A to PIF5 (Fig. 4C, E, F, G and Supplemental Fig. 5) chromatin residence of TOC1 at the *CCA1* promoter is unaffected in a *pif3/4/5* mutant (Supplemental Fig. 6B), similar to results we and others have reported for hypocotyl growth-related genes (*27, 53*). These results indicate that PIF5 is not involved in anchoring or recruiting TOC1 to the *CCA1* promoter.

Notably, ChIP-PCR at the G-box, a DNA motif common to both *CCA1* and *LHY* (Fig 4A), showed a loss of 5X chromatin occupancy only at the *CCA1* promoter, similar to that seen for the *FBS* and *TBS* DNA elements (Supplemental Fig. 4A). This probably resulted from the close proximity of FBS, TBS and G-box sequences (within 140 bp; Fig. 4A) and the low resolution of ChIP-qPCR.

### Absence of FHY3 strongly diminishes TOC1 presence at *CCA1*

To investigate if FHY3 affects chromatin occupancy of TOC1 phospho-variants at *CCA1* and *LHY*, we examined chromatin binding of *TOC1* and 5X in the *fhy3 toc1* background. Absence of FHY3 resulted in 87% lower TOC1 binding to the *CCA1 FBS* site, in contrast to no effect on TOC1 binding to *LHY* promoter. Similar to 5X in the *toc1* single mutant, 5X binding in the absence of FHY3 was also nearly absent at the *CCA1* promoter (Fig. 5A), indicating FHY3 anchors TOC1 to the *CCA1* promoter through the *FBS* element. These results, together with Fig 4E, show that FHY3-TOC1 binding affinity is enhanced by TOC1 phosphorylation, leading to increased presence at the *CCA1* promoter.

**Figure 5.**
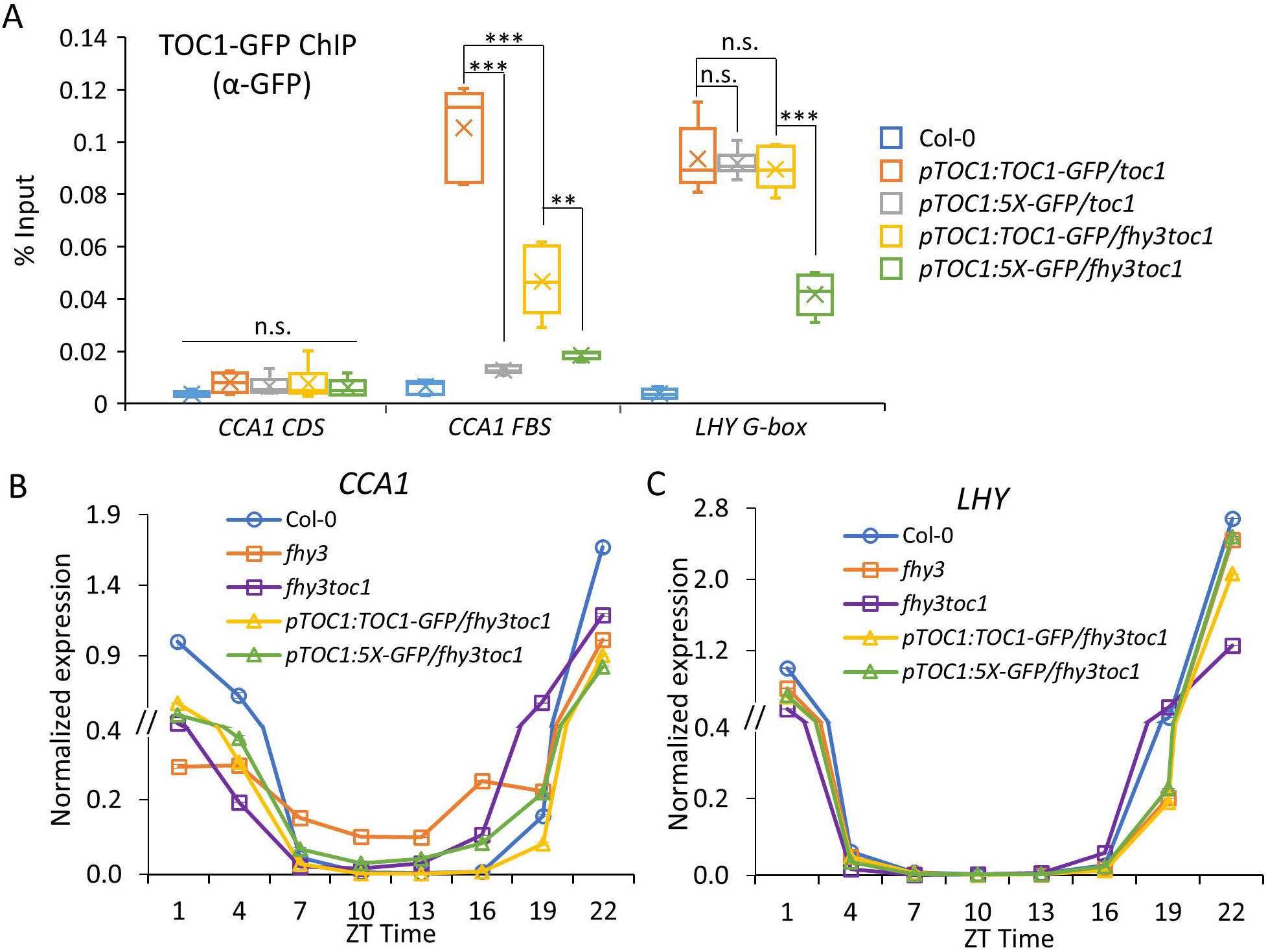
FHY3 is required for TOCl binding to the *CCAJ* promoter and robust *CCAJ* oscillation. **A.** ChIP-qPCR ofTOCl-GFP and 5X-GFP’s residence at target promoters at ZT14 in *tocl andjhy3 tocl* mutant backgrounds. TOCl-GFP was immunoprecipitated by a-GFP with magnetic protein G beads. Data were averaged from 3 independent trials, each trial with 2 technical repeats. Error bars indicate SEM. Asterisks indicate significant differences *(*p* < 0.05, *\*\*p* < 0.001, *\*\*\*p* < 0.0001, one-way ANOVA followed by two-tail Student’s t-test), n.s. = not significant. **B, C.** Time-series expression of *CCAJ* and *LHYin Col-0,jhy3,* wild type and phospho-mutant lines of *TOCJ injhy3 tocl* mutant background. Shown is a representative trial from two independent experiments, error bars indicate SEM. Expression was normalized to Col-0 at ZT1.

Consistent with the absence of *FBS* element in the *LHY* promoter, TOC1 occupancy at *LHY* promoter was the same in the *fhy3 toc1* and *toc1* backgrounds (Fig. 5A). 5X in *fhy3 toc1* showed significantly reduced binding to *LHY* than 5X in *toc1* but this likely resulted from the lower TOC1 protein abundance when *pTOC1:5X-GFP* was crossed into *fhy3 toc1* (Supplemental Fig. 7A).

Together, these results suggest that the selective regulation of *CCA1* by TOC1 phosphorylation results from the anchoring of TOC1 by FHY3 through its *FBS* element, whereas *LHY* lacks *FBS* and TOC1 residence is likely through the G box and independent of its phosphorylation state.

### *fhy3* mutant derepresses *CCA1* expression, similar to *5X* and *S175A*

We next examined the consequences of FHY3 absence and TOC1 phosphorylation on *CCA1*transcript oscillations under light/dark cycles. *fhy3* showed a prominently higher, de-repressed expression of *CCA1* from ZT 4 to 19 (Fig. 5B and Supplemental Fig. 8), consistent with reduced TOC1 presence at *CCA1* in absence of FHY3 (Fig. 3A). The expression phase of *CCA1* in *fhy3 toc1* was slightly earlier than in *fhy3*, like that seen in the *toc1* mutant (Fig. 3B), indicating TOC1 drives phase relationships even in light/dark cycles (Fig. 5B, C and Supplemental Fig. 8). *pTOC1:TOC1-GFP/fhy3 toc1* over-complemented *fhy3 toc1* to a Col-0-like *CCA1* expression pattern possibly due to higher than endogenous TOC1 levels in *pTOC1:TOC1-GFP*; both *pTOC1:TOC1-GFP* lines showed a 1 hour longer period than Col-0 (Fig.1A). *pTOC1:5X-GFP* failed to rescue *toc1* and displayed similar *CCA1* expression pattern as *fhy3 toc1* (Fig. 6B and Supplemental Fig. 8).

**Figure 6.**
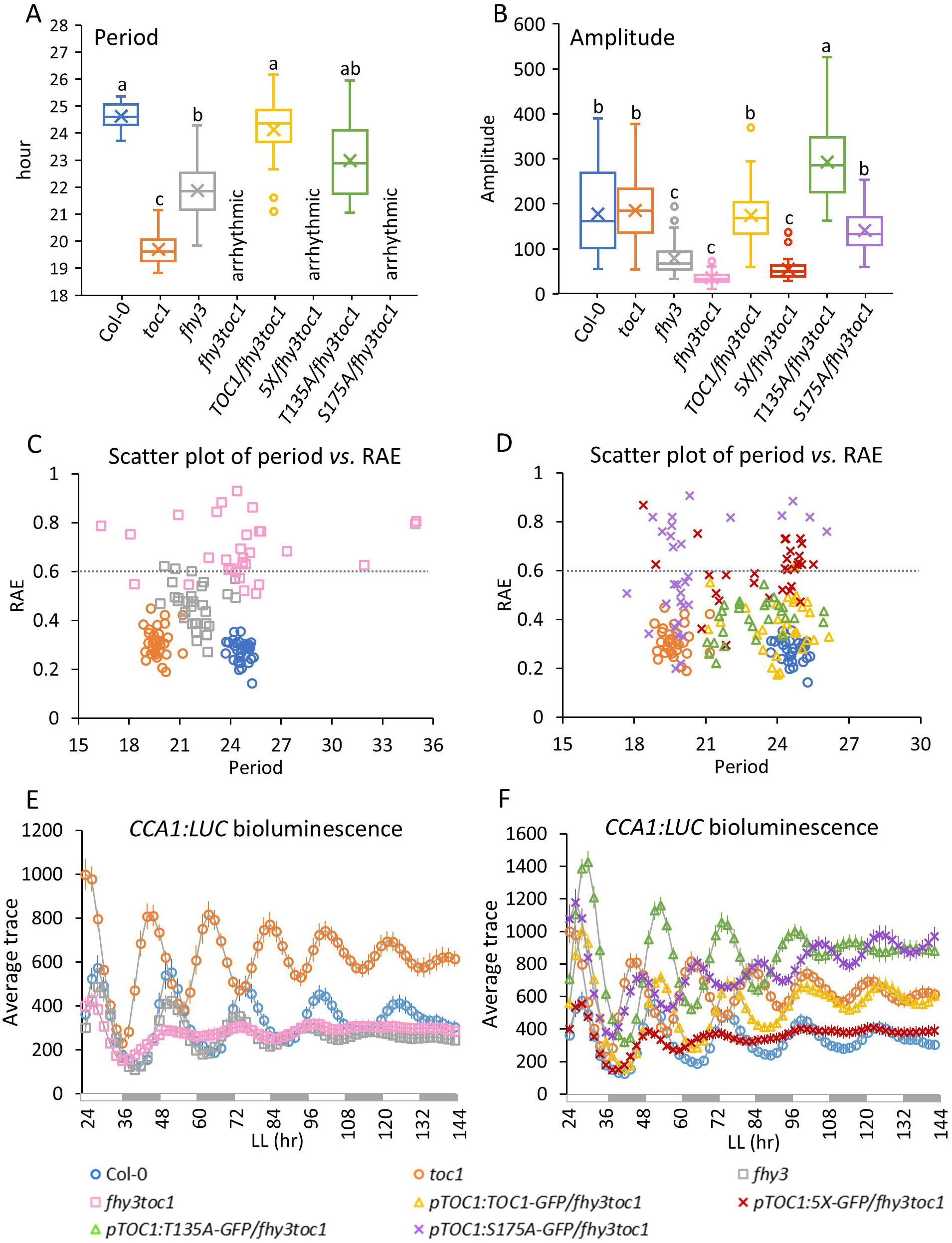
TOCl phosphorylation and FHY3 are required for robust *CCAJ* circadian oscillations. Free­ running period **(A),** amplitude **(B),** RAE **(C, D)** and average *CCAJ-LUC* bioluminescence **(E, F)** of Col-0, *tocl,jhy3,jhy3 tocl* and *TOCJ* native promoter lines *injhy3 tocl* double mutant backgrounds. In A and B, the middle line of the box represents the median, x represents the mean, the bottom line and the top line indicate the 1st and 3rd quartile, respectively. The whiskers extend from the ends of the box to the minimum value and maximum value. Dashed lines in C and D indicate RAE cutoff for the robustness of circadian clock. Seedlings were entrained in 12-h/12-h light/dark cycles for 7 days and then transferred to constant white light at ZT2 for image acquisition at 2-h intervals for 1 week. 30 seedlings from 2 independent trials were averaged. Different letters indicate statistically significant differences *(p* < 0.01, one-way ANOVA followed by Tukey-Kramer HSD test).

In contrast, loss of FHY3 and/or TOC1 phosphorylation did not change *LHY* expression (Fig. 5C and Supplemental Fig. 8). Together with the ChIP-qPCR results, these findings show that FHY3 and TOC1 phosphorylation have no effect on *LHY* transcription due to the absence of *FBS* site on *LHY* promoter.

### FHY3 and TOC1^S175^ phosphorylation are required for *CCA1* rhythmicity

We next examined how phosphorylation affects the coordination between TOC1 and FHY3 in oscillator function. *fhy3* single mutants display a much shorter circadian period (21.9 h) and significantly dampened *CCA1-LUC* oscillation compared to Col-0 under constant white light (Fig. 6A, B and E)(Supplemental Table 2). *CCA1-LUC* oscillations in *fhy3 toc1* double mutants are further dampened and largely arrhythmic (Fig. 6A, B, C and E). In contrast, *fhy3* and *toc1* single mutants displayed RAE within a range (< 0.6) indicative of robust rhythmicity (Fig. 6C). These results show that *FHY3* and *TOC1* act synergistically in sustaining robust circadian period and oscillation.

In contrast, wild-type TOC1 and TOC1^T135A^ over-complemented *fhy3 toc1* and exhibited wild type-like circadian period and amplitude (Fig. 6A and B), probably due the higher than endogenous levels of *TOC1-GFP* expression, consistent with the slightly longer than WT (Col-0) period of *pTOC1:TOC1-GFP*/*toc1* (Fig. 1A). However, *5X* and *S175A* were unable to complement *toc1* in the *fhy3 toc1* mutant, displaying arrhythmic and dampened *CCA1-LUC* phenotype, similar to *fhy3 toc1* (Fig. 6A and F). Furthermore, the oscillations in *5X/ fhy3 toc1* and *S175A/ fhy3 toc1* were much less robust, with half of the seedlings at RAE higher than 0.6 (Fig. 6D). This suggests that TOC1 phosphorylation is required for oscillator function and S175 is the most essential residue.

### Transcriptome comparison between *TOC1* and *5X* lines revealed genome-wide changes in circadian rhythmicity of oscillating genes

The above findings demonstrate the very specific role FHY3 and its partnership with phosphorylated TOC1 plays in the regulation of CCA1 expression. However, to gain a broader insight into the genome-wide effects of TOC1 phosphorylation on the rhythmicity of gene expression, we next performed RNA-seq analyses over a 24-h time course using *pTOC1:TOC1-GFP* and *pTOC1:5X-GFP* lines. The experiments were conducted under 12h L/12h D to better observe the effects of 5X on both phase and amplitude of global gene expression under entrainment, and closer to a natural light/dark environment. In the *TOC1* and *5X* lines similar numbers of expressed genes (23,393 and 23,312, respectively) and rhythmically expressed genes (10,206 [43.6%] and 9,604 [41.2%]) were observed (Fig. 7A)(Supplemental Table 3 and 4).

**Figure 7.**
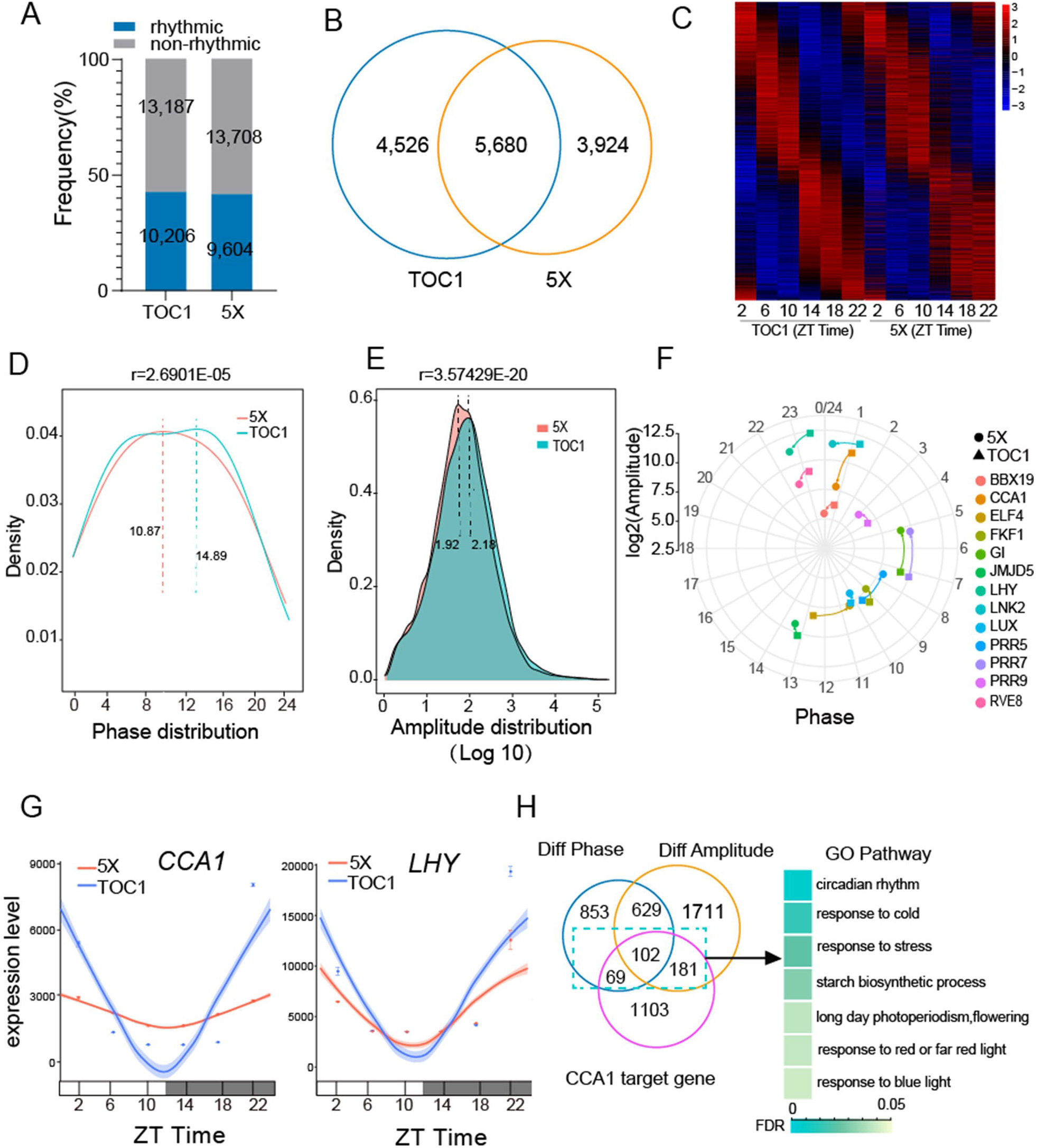
TOCl phosphostate alters phase and amplitude of rhythmic genes. **A.** Number and percentage of rhythmic and non-rhythmic genes in *TOCJ* and *5X* lines identified by time-series RNAseq under 12 h light/ 12 h dark cycles. **B.** Venn diagram showing the overlap of rhythmically expressed genes in *TOCJ* and *5X* lines. **C.** Heatmap of the expression pattern of 5,680 common rhythmic genes sorted by the oscillation phases in *TOCJ* and *5X* lines. **D.** The phase distribution density curves of rhythmic transcripts in *TOCJ* and *5X* plants. The highest density of rhythmic transcripts in *5X* and *TOCJ* plants are phased at 10.87 and 14.89 hrs respectively. **E.** The amplitude density curves of rhythmically expressed genes in *TOCJ* and *5X* lines. The y axis is the log_10_ of the amplitude of gene oscillation calculated by Bio-cycle. The highest density of rhythmic transcripts in *TOCJ* and *5X* plants are with amplitude at I 5I .36 (log_10_ 15I .36=2.18) and 83. I 8 (log_10_ 83.18=1.92), respectively. **F.** Amplitude reduction and phase shift shown in the core clock genes due to impaired phosphorylation of TOCl. The angular coordinates represent phase 0-24h. The log_10_ transformed (peak-trough) represent the amplitude of the gene oscillation and are plotted as radial coordinates. Circles represent 5Xlines and squares represent *TOCJ* lines. **G.** The sinusoidal model fitting traces for the expression patterns of *CCAJ* and *LHY* in *TOCJ vs. 5X* lines that were determined by RNA-seq. The actual data was plotted as mean ± SEM, the confidence interval was plotted as the shade of each curve. **H.** Venn diagram and subsequent GO analysis of *CCA I* target genes (ChIP-seq; Nagel et al., 2015) that have altered phase and/or amplitude (phase change> 2 h; amplitude change> 1.5-fold) in 5X relative to TOCl.

5,680 genes were identified to commonly oscillate in both *TOC1* and *5X* lines, while 4,526 genes and 3,924 genes were oscillating only in *TOC1* or *5X* plants, respectively (Fig. 7B)(Supplemental Table 5). Gene Ontology (GO) terms of circadian rhythm and carbohydrate biosynthetic process were two of the most strongly enriched among the common cycling genes, indicating these processes as continuing to be rhythmic even absent the phosphosites mutated in 5X. TOC1-specific categories of protein glycosylation, cell wall biogenesis and response to stress were specifically strongly enriched, while categories related root development were among the highly enriched *5X*-specific rhythmic genes (Supplemental Fig. 9).

The heatmaps of common rhythmic transcripts between *TOC1* and *5X* revealed differential expression patterns, with the *5X* exhibiting dampened expression peaks in genes phasing at ZT 14 but increased expression peaks in genes phasing at ZT22 (Fig. 7C). The density curves of phase distribution of the common rhythmic genes in *TOC1* and *5X* plants showed a ∼4 hours advanced phase of the most accumulated transcripts in *5X vs*. *TOC1* line (Fig. 7D; Supplemental Fig 10). As indicated with the heatmaps, the *TOC1* line showed the highest density of genes phasing at 14.89, while the *5X* line had the highest number of genes phasing at 10.87 (Fig. 7D). These results are consistent with the shorter period of the 5X line (Fig 1), manifest as a global gene expression phase advance.

Additionally, the amplitude distribution showed a forward shifted peak in the *5X* line indicative of a more dampened gene oscillation in *5X* than in *TOC1* plants. The most enriched rhythmic transcripts in *TOC1* and *5X* plants had amplitude at 151.36 (log_10_ 151.36=2.18) and 83.18 (log_10_ 83.18=1.92), respectively (Fig. 7E).

Further analyses of only core clock genes showed alterations in their rhythmicity even under 12 h light/12 dark cycles, with most genes exhibiting slightly advanced phase and dampened amplitude in the 5X background (Fig. 7F; Supplemental Fig. 11)(Supplemental Table 6). In support of our targeted qPCR results (Fig. 3B-F), genome wide RNA-seq data showed a nearly six-fold reduction in CCA1 amplitude in the 5X background (Fig 7G). In contrast, *LHY* expression did not show a prominent expression difference regardless of TOC1 phosphorylation status (Fig 7H), consistent with our qPCR results (Fig 3B-E). These results further support our notion of the selective regulation of targets through TOC1 interactions with different co-factors.

We next considered that the altered rhythmicity of the cycling genes in 5X plants partially resulted from the altered waveform (i.e. higher trough and earlier phase) of CCA1. When we compared the differentially altered (phase or amplitude differences) oscillating genes in *5X* with known CCA1 target genes (*54*), 352 genes identified as direct targets of CCA1 were enriched (Supplemental Table 7 and 8). These genes are predominately involved in circadian rhythm, cold and stress responses, starch biosynthesis, flowering and light signaling, among which are well-known transcription factors involved in these processes (e.g. *PIF*s, *BBX*s, *PRR*s, *CBF1* and *COR*s) (Fig 7H). Thus, through our analysis of TOC1 phosphorylation state we find support for the notion that CCA1 is both directly and indirectly involved in the transcriptional control of these responses to abiotic and developmental signals.

### TOC1 occupancy at rhythmically expressed genes is enriched by phosphorylation

We next sought to determine whether the rhythmic divergence between TOC1 and 5X can be accounted for by differences in chromatin occupancy at target genes.

Genome-wide ChIP-seq analyses of TOC1 and 5X chromatin binding intensity at ZT14 (the time point of equivalent TOC1 and 5X protein levels) was conducted using *pTOC1:TOC1-GFP* and *pTOC1:5X-GFP* lines. 4846 and 5148 target peaks were identified in *TOC1* and *5X* lines, respectively, of which 3773 were in common (Fig. 8A)(Supplemental Tables 9-11). GO pathway analyses revealed that response to hormone and abiotic stimulus were highly and commonly enriched regardless of TOC1 phosphorylation state (Supplemental Fig. 12C). Of the target genes common between *TOC1* and *5X* , 5X showed similar chromatin binding intensity, but presence of small peaks at gene bodies and distant promoter regions (Fig. 8B) suggests that 5X target binding is less stringent than TOC1.

**Figure 8.**
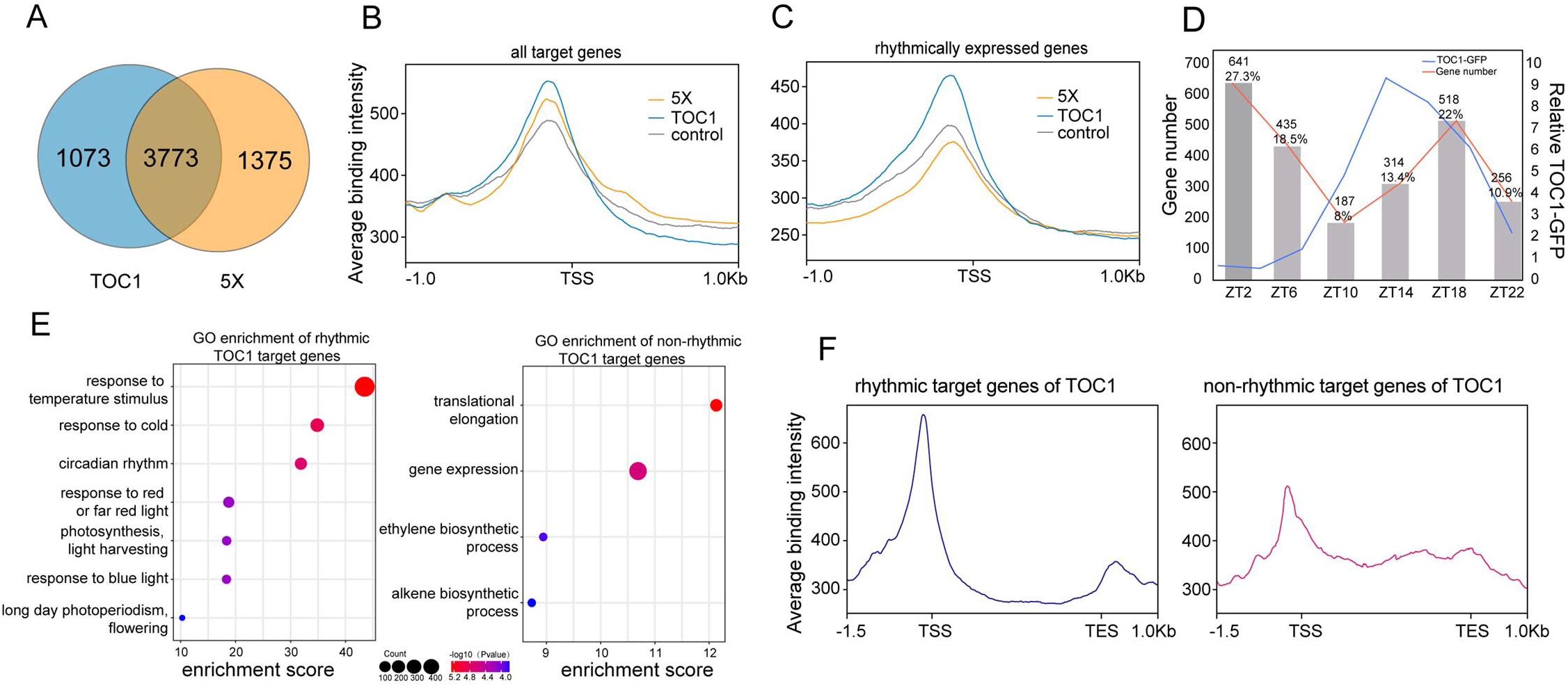
TOCl phosphostate is central to promoter binding to rhythmically expressed genes. **A.** Venn diagram showing the overlap of targets in *TOCJ* and *5X* lines. **B, C.** Binding profiles of TOCl and 5X at the TSS ± 1kb on all target genes **(B)** and rhythmic target genes **(C)** at ZT14. TOCl binding profile at ZT2 was used as control. **D.** Quantitative profiling of rhythmic TOCl target genes identified through time-course RNA-seq. Genes exhibiting rhythmic expression patterns (p<0.05) were classified into six phase-specific clusters using the Bio-Cycle algorithm, with cluster distribution frequencies annotated above corresponding histogram bars. Relative TOC1-GFP protein abundance driven by native TOCl promoter is also shown demonstrating anti-phase oscillation relative to gene cluster distribution of TOCl target genes (red line). **(E)** GO analysis of the rhythmic and non-rhythmic genes bound by TOCl. The enrichment score is the negative natural logarithm of the enrichment P-value derived from the Fisher’s exact test. Count indicates the number of genes has been identified in each pathway. **(F)**

More notably, examination of the chromatin binding profile using only rhythmically expressed target genes shows the intensity of 5X was prominently lower than TOC1 (Fig. 8C), suggesting that phosphorylation of TOC1 promotes chromatin presence more robustly on rhythmic genes.

To better compare the nature of rhythmic TOC1 target genes with non-cycling TOC1 target genes we first clustered the cycling genes by time of day (phase) (Supplemental Fig 13) and then plotted them across six phase-specific time points superimposed with TOC1 protein abundance (Fig 8D). Peak phase expression of the TOC1 target genes is generally anti-phase to TOC1 protein abundance, supporting previous evidence that TOC1 acts as a transcriptional repressor.

Divergence from a perfectly anti-phase relationship between TOC1 abundance and peak phase of gene expression may arise from TOC1 co-factors modifying peak TOC1 efficacy.

Comparative GO analysis between cycling and non-cycling TOC1 genes showed a strong divergence in the nature of the targets (Fig 8D). Unsurprisingly, circadian rhythm and response to temperature and cold were the most strongly enriched cycling TOC1 target genes. In contrast, general gene expression and genes involved in translational elongation were the most commonly enriched non-cycling TOC1 target genes (Fig. 8E). However, the TOC1 binding profile of the non-cycling genes showed much greater presence of TOC1 throughout the gene body (Fig 8F). This suggests that some non-cycling gene residence of TOC1 may not be functionally significant.

### FHY3 and TOC1p shape the rhythmicity of the cyclic genes involved in circadian clock, light responses and photoperiodism

Consistent with our ChIP-qPCR results (Fig. 2A), our genome-wide ChIP-seq results confirm that 5X is nearly absent at the *CCA1* promoter, resulting in significantly elevated CCA1 expression at ZT14 (Fig. 9A). In contrast, *LHY* is similarly bound by TOC1 and 5X, reflecting a similarly repressed expression at night (Fig. 9A; Fig 2A). Other core clock genes targeted by TOC1, such as PRR9, PRR7, GI and ELF4, are bound more weakly (∼30%) by 5X than TOC1, similar to our ChIP-qPCR results (Fig. 2A), leading to enhanced expression of these genes in *5X* plants (Supplemental Fig. 14). These results show that phosphorylation is crucial to TOC1 chromatin binding specificity and strength, particularly on cycling genes, leading to altered rhythmicity of their expression.

**Figure 9.**
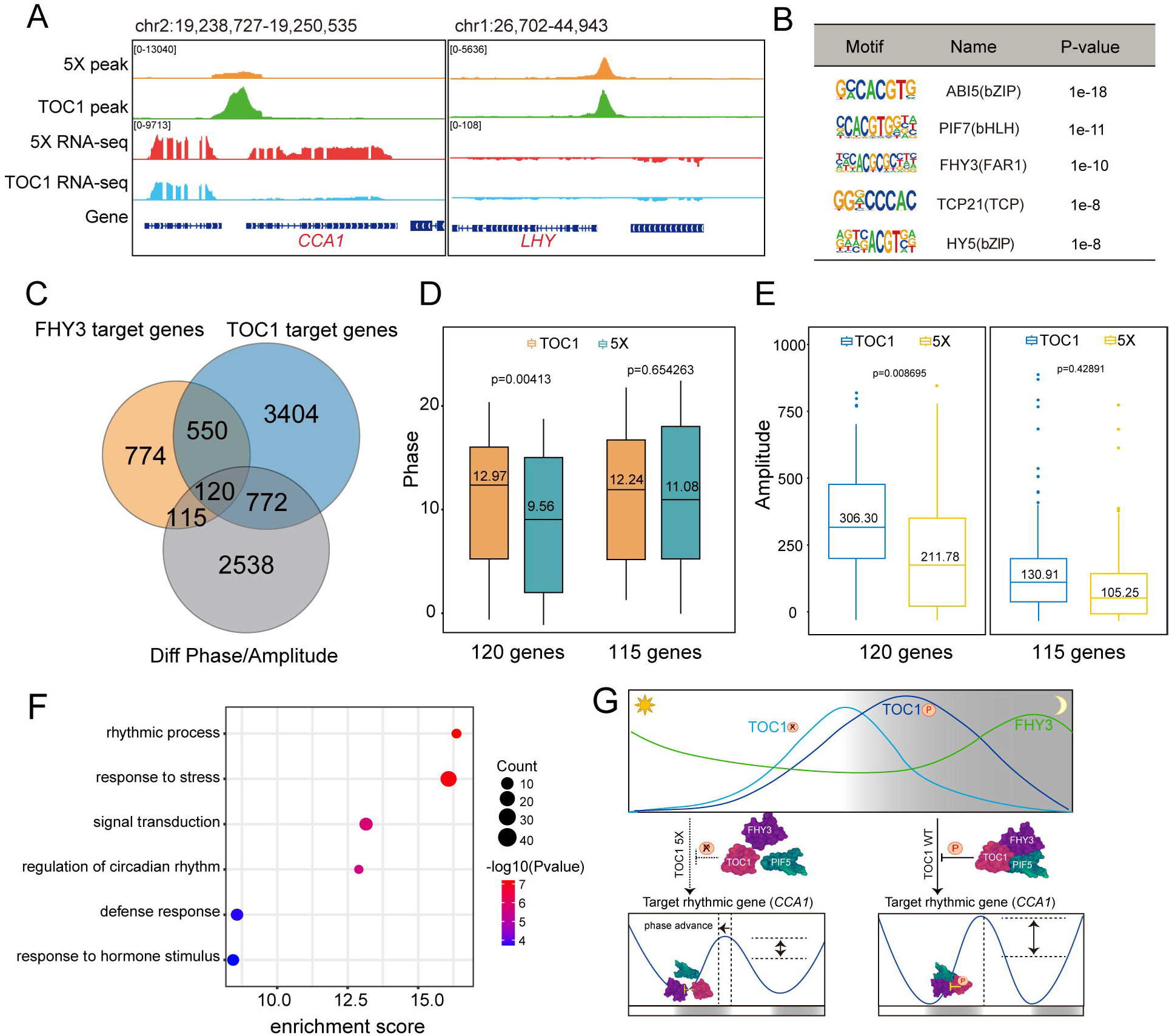
FHY3 and TOCl phosphostate shape robustness and period of rhythmically expressed genes. . **A.** Integrative Genomics Viewer (IGV) view of ChIP-seq profiles and RNA-seq data at *CCAJ* and *LHYlocus* in *TOCJ* and 5Xlines. The yellow and green wiggle plot represents the peak of ChIP-seq track. The red and blue wiggle plots represent the peak of RNA-seq track. Up (CCAl) and down (LHY) trace in RNA-seq track indicate forward- and reverse-strand RNA reads, respectively. **B.** The position weight matrix (PWM) of motifs enriched in differentially oscillating genes bound by TOCl, along with their optimal enrichment p-values and candidate TFs shown in the table. **C.** Venn diagram showing the overlap of differentially oscillating genes bound by FHY3 and TOCl. **D, E.** Comparison of the phase **(D)** and amplitude **(E)** of differentially oscillating transcripts (phase change> 2 h; amplitude change> 1.5-fold) bound by both FHY3 and TOCl (120 genes) or only by FHY3 (115 genes) between the *TOCJ* and *5X* ChIP-seq datasets. **F.** GO analysis of the 120 differentially oscillating genes bound by FHY3/TOC1. The enrichment score is the negative natural logarithm of the enrichment P-value derived from the Fisher’s exact test. Count indicates the number of genes has been identified in each pathway. **G.** Proposed mechanism for the role of TOC1 phosphorylation in the control of CCAl gene expression. During the day, early-phased SX (unphosphorylated TOCl) increases faster than wild-type TOCl (TOClWT), resulting in stronger inhibition of FHY3 despite the poorer interaction with FHY3, resulting in early reduced CCAl transcription. At night, SX levels decreases more rapidly and interacts more poorly with FHY3 than wild-type TOCl, releasing FHY3 for CCAl activation which raises the trough of CCAl. The TOC1-PIF5 interaction, also phosphostate dependent, act together to repress FHY3 activity. Not shown are the NF-YB and C co-factors that complex with TOCl at the chromatin.

When TOC1 target genes from our ChIP-seq are compared to the targets identified from Huang et al. (*48*) we identified 64.2% of the target genes from the Huang et al. study, despite different TOC1 transgenic lines and p-value cutoffs used for the two datasets. This comparison indicates a good degree of consistency of our ChIP-seq with previous reports (Supplemental Fig. 12A). Motif analysis of TOC1-bound sequences identified significantly enriched G-Box that are bound by the basic helix-loop-helix (bHLH) and basic Leu zipper (bZIP) TFs, such as ABFs and PIFs (Supplemental Fig. 12B). HY5, FHY3 and TCP binding motifs were also identified among TOC1 targeting sequences (Fig. 9B), consistent with our prior ChIP-qPCR results.

To identify potential transcriptional co-regulators of TOC1 target genes and the effect of phosphorylation, we compared oscillating genes showing altered phase or amplitude in *5X* lines with genes bound differentially by 5X. Motif analysis of the overlapped genes identified the FHY3 binding element among those significantly enriched in the rhythmic genes bound by TOC1 (Fig. 9B), consistent with the essential role of FHY3 in maintaining the robustness of circadian rhythms together with TOC1. To identify additional genes that are co-regulated by TOC1 and FHY3, we overlapped the phase and amplitude-altered genes from the 5X data set with genes from FHY3 (*55*) and TOC1 ChIP-seq datasets. From this three-way comparison 120 genes altered in 5X plants were bound by both FHY3 and TOC1 (Supplemental Table 12), while 115 genes were bound only by FHY3 (Supplemental Table 13)(Fig. 9C). The 120 overlapping genes were phase advanced by more than 3 hours and the amplitude of these genes decreased by ca. 31 % in *5X* plants (Fig. 9D, E). In contrast, the 115 genes that were only bound by FHY3 exhibited much reduced amplitude compared to the 120 genes, with similar phase and amplitude between *TOC1* and *5X* plants (Fig. 9D, E). The 120 overlapped genes were primarily enriched for GO terms describing rhythmic processes, response to stress, signal transduction and regulation of circadian rhythm, consistent with known targets of TOC1 and FHY3 (Fig. 9F). However, for the 115 non-overlapped genes, no GO terms were enriched. Together, these results suggest that FHY3 and phosphorylated TOC1 are essential to the general maintenance of robust gene oscillations, as we have specifically shown here for *CCA1*.

## DISCUSSION

### FHY3 is the key co-factor interacting with TOC1 to selectively regulate *CCA1* expression

TOC1 is a core transcriptional repressor in the plant circadian clock. With peak expression in the early evening, genome wide studies have shown a wide-ranging effect of TOC1 on gene expression well beyond the core elements of the circadian oscillator (*13, 48, 56*)(this study). Additionally, although TOC1 is phosphorylated (*24, 27*), the role of this modification in the context of the circadian system has been unclear. Here we have shown that TOC1 as a repressor of FHY3 transcriptional activation of *CCA1* depends on the phosphorylation of N-terminal residues, particularly serine 175.

The failure of the S175A mutation to rescue the short period *toc1-1* mutation (Fig. 1) suggests a deficiency in the transcriptional repressor functions of TOC1^S175A^. We show that two aspects of TOC1 function in the circadian system are affected by phosphorylation: protein-protein interactions and chromatin residence. TOC1 binds DNA via the carboxy-terminal CCT domain (*57*), but requires NF-YB and NF-YC cofactors for robust presence at the promoters of hypocotyl growth-related genes, and N-terminal TOC1 phosphorylation to promote formation of the NF-TOC1 trimeric complex (*27*). Here we have extended these findings to show that full TOC1 chromatin binding at core circadian clock genes similarly requires members of the NF-YC family and these same phosphosites (Fig. 2).

The phosphostate-dependent binding of TOC1 to the *CCA1* promoter (Fig. 2A) prompted a more detailed investigation into the cause. Using the relative insensitivity of the TOC1 phosphostate to *LHY* promoter binding for comparison, we identified the FHY3-TOC1 protein interaction as the primary determinant of the TOC1 phosphostate-dependent difference between the *CCA1* and *LHY TOC1* chromatin presence, and the cause for the increased *CCA1* expression in the 5X, S175A and *fhy3* mutant backgrounds (Fig. 3). Previous work established TOC1 amino acid residues 141-618 as necessary for FHY3 interaction (*18*). Within this region are 4 of 5 serine/threonine residues mutated in 5X (T135, S175, S194, S201, S204) including the single most effective mutation, S175A. Alanine substitution of T135 that was not within the interaction region exhibited trivial changes in the circadian rhythm, FHY3 interactions and chromatin residence (Figs. 1, 3 and 4). Our findings now confirm and expand on that work to show that phosphorylation of TOC1 at S175 is crucial to the interaction.

Interestingly, previous work has implicated both FAR1 and FHY3 as important regulators of circadian function (*18, 58*). However, roles for these two closely related transcription factors in the activation of *ELF4* and *CCA1/LHY* through promoter binding has tended to emphasize FHY3 as the more important of the pair. In most cases *fhy3* mutants are more severe than *far1* lines, with the double mutant often additive. Here we show a strong difference in the relative roles of the two factors in how TOC1 controls *CCA1* expression, consistent with these previous reports. While other work has shown a TOC1-FAR1 interaction in yeast two hybrid and bifluorescence complementation assays (*18*), we were unable to demonstrate measurable interactions. Our *in planta* co-IPs between TOC1 and FAR1 showed very weak to no interaction, in strong contrast with the significant phospho-state dependent interaction between TOC1 and FHY3 (Fig 4; Supplemental Fig 5). Different assays and experimental conditions may account for the different results, but our findings are consistent with most reports emphasizing the primary role for FHY3 over FAR1 in light signaling and circadian systems (*18, 59, 60*).

PIF5 has been implicated to act with TOC1 to repress FHY3 activation of *CCA1* transcription (*18*). Our results show TOC1 binding to *CCA1* and *LHY* promoters is independent of PIF5 presence, although TOC1 interaction with PIF5 is phosphostate-dependent (Supplemental Fig. 6B, Fig 4E). PIF5, and PIFs in general, are associated with circadian regulation (*61*). *pif5* single mutants have no effect on *CCA1* expression, although PIF5 can be found at the *CCA1* promoter and strong ectopic PIF5 expression can repress *CCA1* expression (*18*). Only combinatorial stacking of multiple *pif* mutants alters circadian period, suggesting genetic redundancy among the PIF family (*61*). Together with our findings, these results suggest that phosphorylated TOC1 is important in the recruit of one or more PIF species to the FBS/G-BOX vicinity of the *CCA1* promoter to control phase and amplitude under light/dark cycles (Fig. 8J). The role of PIF5 may be related to sucrose sensing, as Shor et al. (2017) showed enhanced binding to the *CCA1* and *LHY* promoters at higher sucrose levels. A PIF5-TOC1 complex might then be a point of confluence where circadian and metabolic signals converge to regulate *CCA1* and *LHY* expression.

*LHY* transcription is repressed by TOC1 (*13, 48*), consistent with TOC1 chromatin presence (Fig. 2 and 3). *In vitro* binding of an NF-TOC1 trimer at the *LHY* promoter supports our findings (*57*). The *toc1* null shows an increase in LHY mRNA during the time of maximum TOC1 protein presence (Fig.3C) (*24*), but the waveform of the mutant is unaffected by the presence or absence (*fhy3 toc1*) of FHY3 (Fig. 5C). Hence, the NF-TOC1 trimeric complex (*27*) is either acting alone (possibly with a histone deacetylase, like HDA15; (*27*)) or with one or more unknown cofactors to repress *LHY* expression via the G-box or related sequences. The additional FHY3-dependent component of *CCA1* regulation, not present for *LHY*, raises the interesting question of what FHY3 contributes to *CCA1* expression, particularly since its phase of circadian expression is so similar to *LHY*. Additionally, paradoxically the transcriptional activator FHY3 is also necessary for TOC1 recruitment to the promoter (Fig. 5), which then acts to repress *CCA1* expression.

### Global rhythmicity changes in gene expression are subject to TOC1 phosphorylation status

Both in circadian studies and in general gene regulation, protein phosphorylation plays a major role determining genome-wide gene expression. (*62–67*). To further understand the importance of TOC1 phosphorylation on a broader scale we examined the genome-wide effects of the 5X mutant on rhythmicity, phase and amplitude of gene oscillations under the more natural conditions of light/dark cycles. The advanced phase of output genes (Fig. 7D) and core clock genes (Fig. 7F) expressed in the 5X background is consistent with the shorter period (Fig. 1) but the reduced amplitude in that background (Fig. 7E) suggests a weakening in the robustness of rhythmic gene expression in general. In some circadian systems, phosphorylation of key clock proteins alters stability, leading to weakened rhythmicity (*66*). 5X and TOC1 protein oscillations were similarly robust under light/dark cycles (Supplemental Fig. 2) and a change in protein abundance alone likely cannot account for lower amplitude of target genes. However, weaker homodimerization and weaker interaction with NF-YB and C components (*27*), leading to poor chromatin binding and reduced gene repression, may be the most likely reason for low amplitude gene expression oscillations in the 5X background. Focusing on 5X effects on CCA1 target genes alone (Fig. 7G), circadian rhythm genes are disproportionally affected (Fig. 8C; Supplemental Fig. 11) which helps explain the overall effects on oscillating genes.

### TOC1 phosphostate disproportionally affects chromatin binding at promoters of rhythmic genes

As a transcriptional repressor, if phosphorylation of TOC1 alters chromatin presence it might be expected to affect binding at all target genes. However, only rhythmically expressed genes showed a much lower binding profile for 5X (compare Fig. 8B and C), consistent with lower amplitude oscillations of the rhythmic genes in this background. This interesting result suggests that a range of different TOC1 cofactors, not just FHY3, contribute to TOC1’s role in sustaining oscillations, and also require TOC1 phosphorylation for optimal functionality.

A similar enrichment of a core circadian transcriptional regulator in Drosophila, CLOCK (CLK), specifically at the promoters of rhythmic genes has been described previously (*68*). As in our study, these researchers also found CLK binding to non-cycling target genes. In our work, non-rhythmic TOC1 target genes (Supplemental Fig 13B) were enriched for translational elongation (Fig 8E), part of the complex process of mRNA translation that has previously been found to be circadian regulated at multiple steps, including initiation, elongation and phase separation (*69*). It is possible that the arrhythmicity or low rhythmicity of these gene transcripts, despite TOC1 binding, is due to their high mRNA stability.

Alternatively, other transcriptional activators binding throughout the diurnal time course could override any phase-dependent effects of TOC1. As well, the greater degree of gene body binding of TOC1 on the non-rhythmic targets (Fig. 8F) may suggest a non-productive presence of TOC1 at some of these genes. Future work will be necessary to distinguish between these possibilities.

## Author contributions

JY and XL designed the methodology, JY and YZ performed experiments with assistance from CG, ZT and DC. GC performed the genomic data analysis. DES conceptualized, supervised the project and wrote the manuscript with JY and comments from XL.

## Data and materials availability

All data needed to evaluate the conclusions in the paper are present in the paper and/or the Supplementary Materials.

