## Supplemental Figures 1-14 for "TOC1 phosphorylation disproportionally enhances chromatin binding at rhythmic gene promoters"

### Slide 1
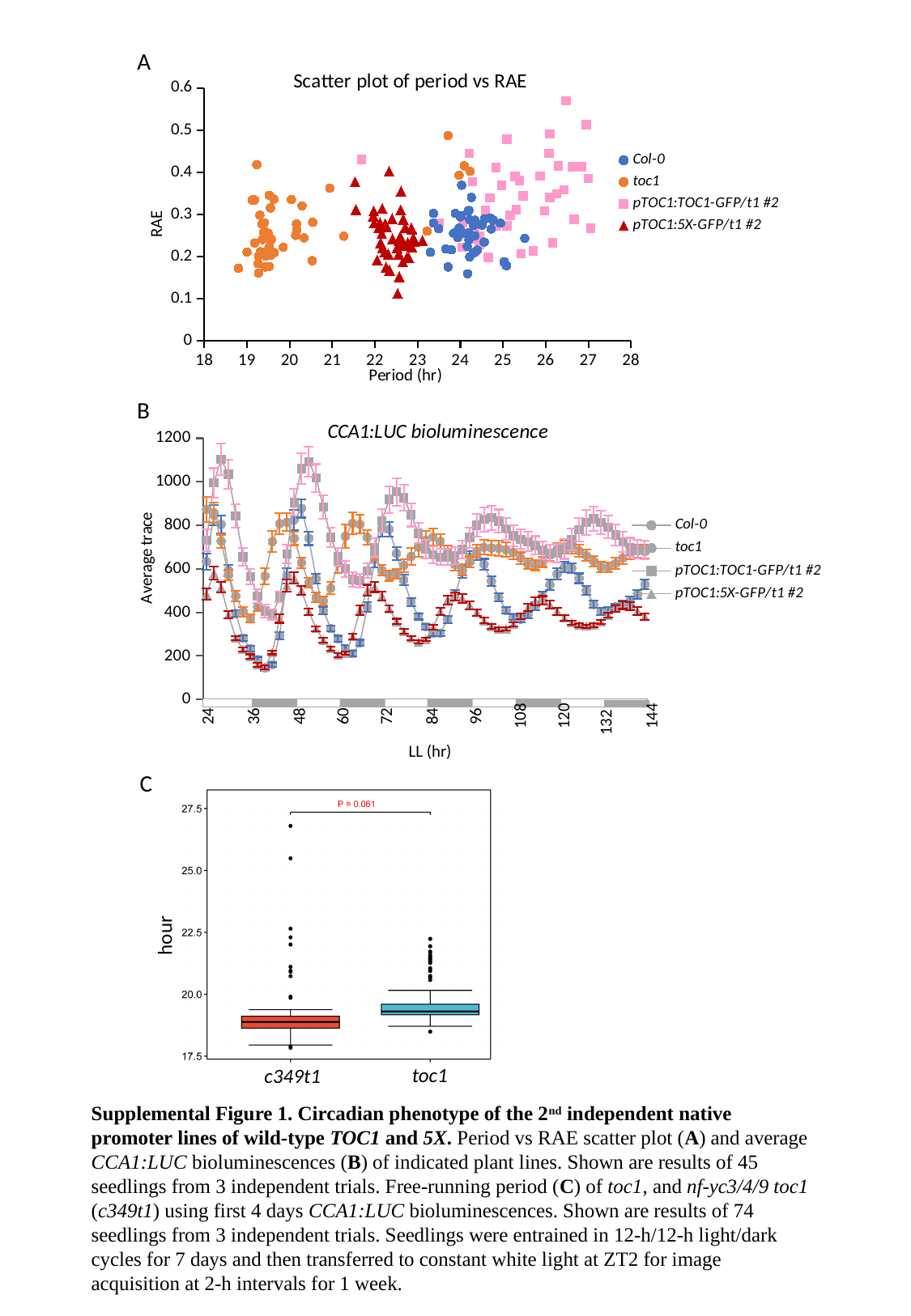

A
#### Chart: Scatter plot of period vs RAE
| Category | Col-0 | toc1 | pTOC1:TOC1-GFP/t1 #2 | pTOC1:5X-GFP/t1 #2 |
|---|---|---|---|---|B
#### Chart: CCA1:LUC bioluminescence
| Category | Col-0 | toc1 | pTOC1:TOC1-GFP/t1 #2 | pTOC1:5X-GFP/t1 #2 |
|---|---|---|---|---|
| 24 | 633.4359259259259 | 873.4542307692308 | 731.834074074074 | 484.26846153846157 |
| 26 | 847.2725925925928 | 858.245 | 994.633333333333 | 580.8653846153846 |
| 28 | 803.6207407407409 | 726.9946153846154 | 1103.5166666666667 | 515.256923076923 |
| 30 | 593.055185185185 | 571.3542307692308 | 1034.4525925925925 | 389.3542307692308 |
| 32 | 394.43407407407403 | 474.9911538461538 | 842.521111111111 | 282.0561538461538 |
| 34 | 282.6196296296296 | 403.84999999999997 | 655.2222222222221 | 228.71 |
| 36 | 233.02740740740742 | 371.71999999999997 | 564.8744444444443 | 196.84403846153845 |
| 38 | 182.4466666666666 | 424.9776923076922 | 475.53333333333325 | 160.35815384615384 |
| 40 | 145.31444444444443 | 566.4353846153846 | 406.2374074074073 | 147.79134615384615 |
| 42 | 159.7998888888889 | 725.5238461538463 | 390.99185185185183 | 215.30461538461537 |
| 44 | 293.8140740740741 | 807.623076923077 | 471.0885185185184 | 371.61730769230775 |
| 46 | 568.0392592592592 | 814.9073076923077 | 669.2166666666666 | 523.895 |
| 48 | 824.9840740740741 | 739.7261538461537 | 904.6977777777778 | 558.7711538461541 |
| 50 | 877.5759259259257 | 628.1453846153846 | 1060.53 | 499.9407692307692 |
| 52 | 740.245925925926 | 536.0561538461538 | 1092.2181481481477 | 403.50615384615384 |
| 54 | 556.1177777777776 | 468.4511538461539 | 1016.6862962962963 | 325.50307692307695 |
| 56 | 407.91481481481486 | 452.48769230769227 | 883.8381481481482 | 272.29307692307685 |
| 58 | 325.76407407407413 | 510.7592307692308 | 744.8970370370371 | 234.37538461538458 |
| 60 | 279.0596296296297 | 626.5503846153845 | 655.8651851851852 | 204.25115384615387 |
| 62 | 234.26962962962963 | 749.4715384615384 | 600.5411111111111 | 212.3161538461539 |
| 64 | 210.23859259259262 | 808.9496153846153 | 552.0103703703703 | 287.8765384615384 |
| 66 | 260.82592592592596 | 804.4215384615386 | 545.0988888888888 | 410.3384615384615 |
| 68 | 423.67333333333335 | 744.7015384615383 | 591.0355555555556 | 501.1580769230768 |
| 70 | 641.7918518518519 | 666.9150000000001 | 698.1229629629629 | 517.121923076923 |
| 72 | 783.132222222222 | 593.825 | 819.9033333333333 | 475.565 |
| 74 | 781.8566666666667 | 569.208076923077 | 918.851851851852 | 417.43730769230774 |
| 76 | 671.3625925925926 | 576.7796153846155 | 953.5548148148149 | 358.1584615384615 |
| 78 | 548.3025925925926 | 615.6773076923077 | 926.4859259259259 | 313.65 |
| 80 | 447.7481481481481 | 657.4265384615386 | 848.3196296296296 | 280.70769230769235 |
| 82 | 381.59333333333336 | 699.9434615384617 | 764.9122222222222 | 264.4242307692308 |
| 84 | 334.66703703703706 | 731.3419230769232 | 691.6529629629629 | 276.2773076923077 |
| 86 | 304.1392592592592 | 745.0896153846153 | 664.3118518518518 | 331.3042307692307 |
| 88 | 304.8862962962962 | 726.0207692307692 | 654.1525925925924 | 403.7719230769231 |
| 90 | 367.4774074074074 | 676.023076923077 | 654.1003703703703 | 457.18153846153837 |
| 92 | 478.65037037037035 | 619.8126923076923 | 661.1659259259259 | 472.18269230769226 |
| 94 | 587.1911111111112 | 596.686923076923 | 688.4618518518519 | 464.16692307692307 |
| 96 | 654.1811111111111 | 631.6923076923077 | 743.7155555555556 | 433.01115384615395 |
| 98 | 668.7655555555555 | 681.1107692307692 | 800.4622222222222 | 399.57923076923083 |
| 100 | 620.3555555555555 | 699.2111538461538 | 827.8066666666665 | 361.87423076923073 |
| 102 | 543.9433333333333 | 695.9269230769231 | 835.1096296296297 | 335.9396153846154 |
| 104 | 468.78629629629626 | 694.8134615384616 | 819.3592592592593 | 322.86 |
| 106 | 408.8359259259259 | 687.9192307692308 | 784.4770370370371 | 324.7853846153846 |
| 108 | 377.31 | 674.801923076923 | 751.9492592592592 | 344.27884615384625 |
| 110 | 369.14444444444445 | 649.3288461538461 | 736.1122222222222 | 382.10653846153855 |
| 112 | 388.8340740740742 | 622.8480769230769 | 725.0433333333333 | 423.27461538461546 |
| 114 | 427.00518518518516 | 616.5438461538462 | 707.7188888888891 | 452.24115384615385 |
| 116 | 472.67999999999995 | 632.3180769230769 | 685.4374074074076 | 455.18500000000006 |
| 118 | 526.3044444444445 | 663.3407692307693 | 671.9633333333334 | 435.20346153846157 |
| 120 | 577.6237037037035 | 689.3873076923077 | 672.1918518518519 | 405.2515384615384 |
| 122 | 609.2314814814815 | 700.8538461538463 | 691.7181481481483 | 373.4100000000001 |
| 124 | 603.1574074074074 | 698.8396153846153 | 734.2033333333333 | 351.60153846153844 |
| 126 | 556.8455555555554 | 683.2819230769231 | 779.3366666666667 | 340.46730769230777 |
| 128 | 500.8381481481482 | 661.1165384615384 | 813.801111111111 | 336.296923076923 |
| 130 | 436.28666666666663 | 634.926153846154 | 829.8870370370373 | 340.74115384615385 |
| 132 | 404.2170370370371 | 608.936923076923 | 813.5322222222222 | 355.5153846153846 |
| 134 | 408.8496296296296 | 605.7065384615385 | 792.114074074074 | 387.39153846153846 |
| 136 | 424.88962962962967 | 625.4703846153845 | 755.122222222222 | 418.0265384615385 |
| 138 | 437.8040740740741 | 649.48 | 723.4785185185186 | 432.3276923076923 |
| 140 | 452.35814814814813 | 672.2034615384616 | 698.8422222222223 | 428.7565384615384 |
| 142 | 481.5292592592593 | 681.5896153846153 | 691.0099999999999 | 406.02384615384614 |
| 144 | 529.0814814814813 | 679.4003846153847 | 686.9944444444445 | 381.41269230769234 |144
108
120
24
36
48
60
72
84
96
132
LL (hr)
C
hour
toc1
c349t1
Supplemental Figure 1. Circadian phenotype of the 2^nd^ independent native promoter lines of wild-type TOC1 and 5X. Period vs RAE scatter plot (A) and average CCA1:LUC bioluminescences (B) of indicated plant lines. Shown are results of 45 seedlings from 3 independent trials. Free-running period (C) of toc1, and nf-yc3/4/9 toc1 (c349t1) using first 4 days CCA1:LUC bioluminescences. Shown are results of 74 seedlings from 3 independent trials. Seedlings were entrained in 12-h/12-h light/dark cycles for 7 days and then transferred to constant white light at ZT2 for image acquisition at 2-h intervals for 1 week.

### Slide 2
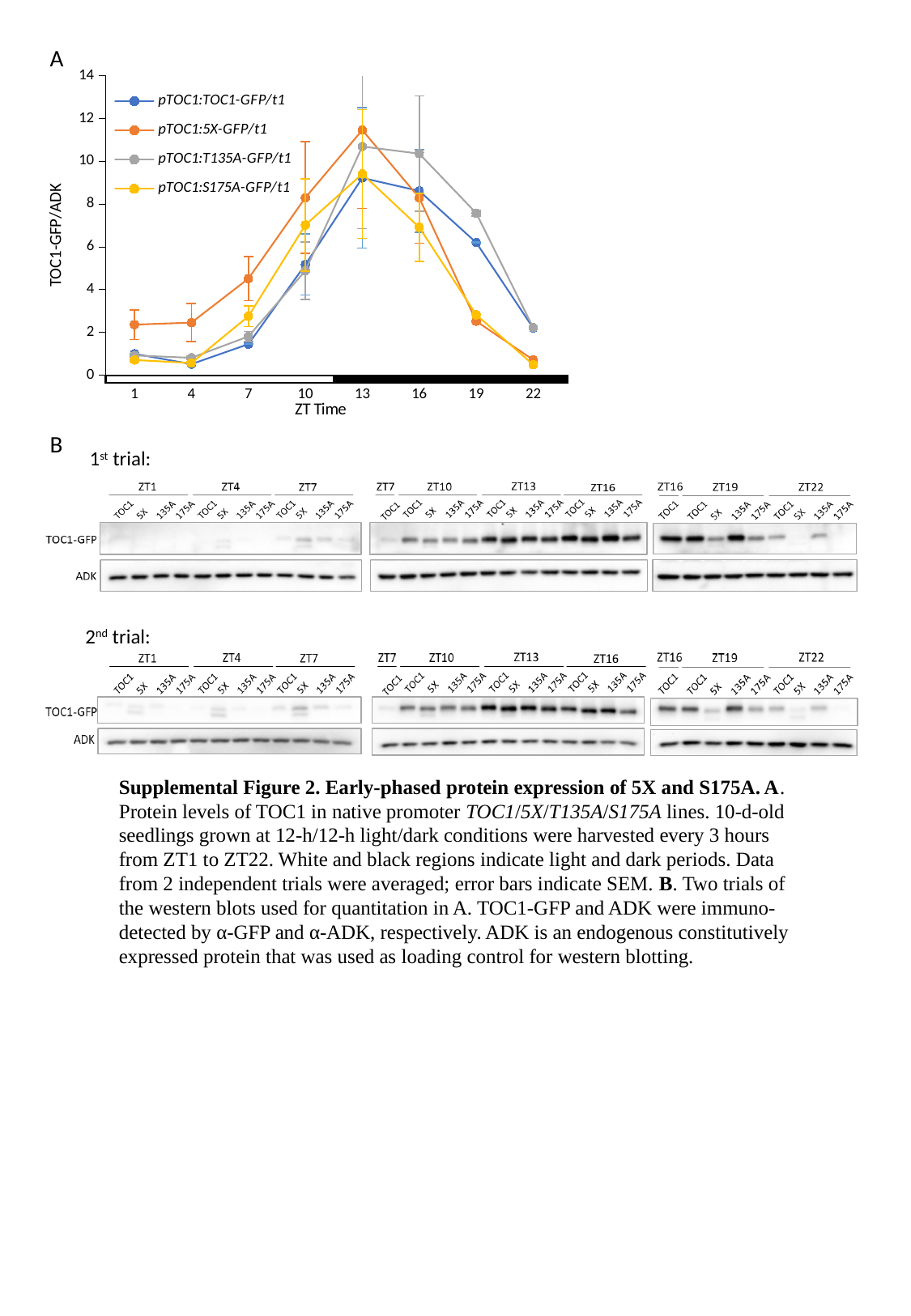

A
#### Chart
| Category | pTOC1:TOC1-GFP/t1 | pTOC1:5X-GFP/t1 | pTOC1:T135A-GFP/t1 | pTOC1:S175A-GFP/t1 |
|---|---|---|---|---|
| 1 | 1.0 | 2.366603035397198 | 0.9266293460437502 | 0.7121699478020826 |
| 4 | 0.5188940509540522 | 2.461230053946285 | 0.8148107172389132 | 0.5707429526859075 |
| 7 | 1.4535928346995277 | 4.518283579422434 | 1.811985976124944 | 2.757328610761653 |
| 10 | 5.172370942649607 | 8.301674586648815 | 4.888622963153501 | 7.022633320512902 |
| 13 | 9.225294180895457 | 11.462719377252721 | 10.695486581291817 | 9.412292343278217 |
| 16 | 8.614588160450438 | 8.292920496010288 | 10.358277961644397 | 6.914905178959699 |
| 19 | 6.199736770857255 | 2.5264957572966362 | 7.575346846809181 | 2.827898006771309 |
| 22 | 2.2055402843158967 | 0.719292028522146 | 2.226725678358838 | 0.4916723807360714 |
B
1^st^ trial:
2^nd^ trial:
Supplemental Figure 2. Early-phased protein expression of 5X and S175A. A. Protein levels of TOC1 in native promoter TOC1/5X/T135A/S175A lines. 10-d-old seedlings grown at 12-h/12-h light/dark conditions were harvested every 3 hours from ZT1 to ZT22. White and black regions indicate light and dark periods. Data from 2 independent trials were averaged; error bars indicate SEM. B. Two trials of the western blots used for quantitation in A. TOC1-GFP and ADK were immuno-detected by α-GFP and α-ADK, respectively. ADK is an endogenous constitutively expressed protein that was used as loading control for western blotting.

### Slide 3
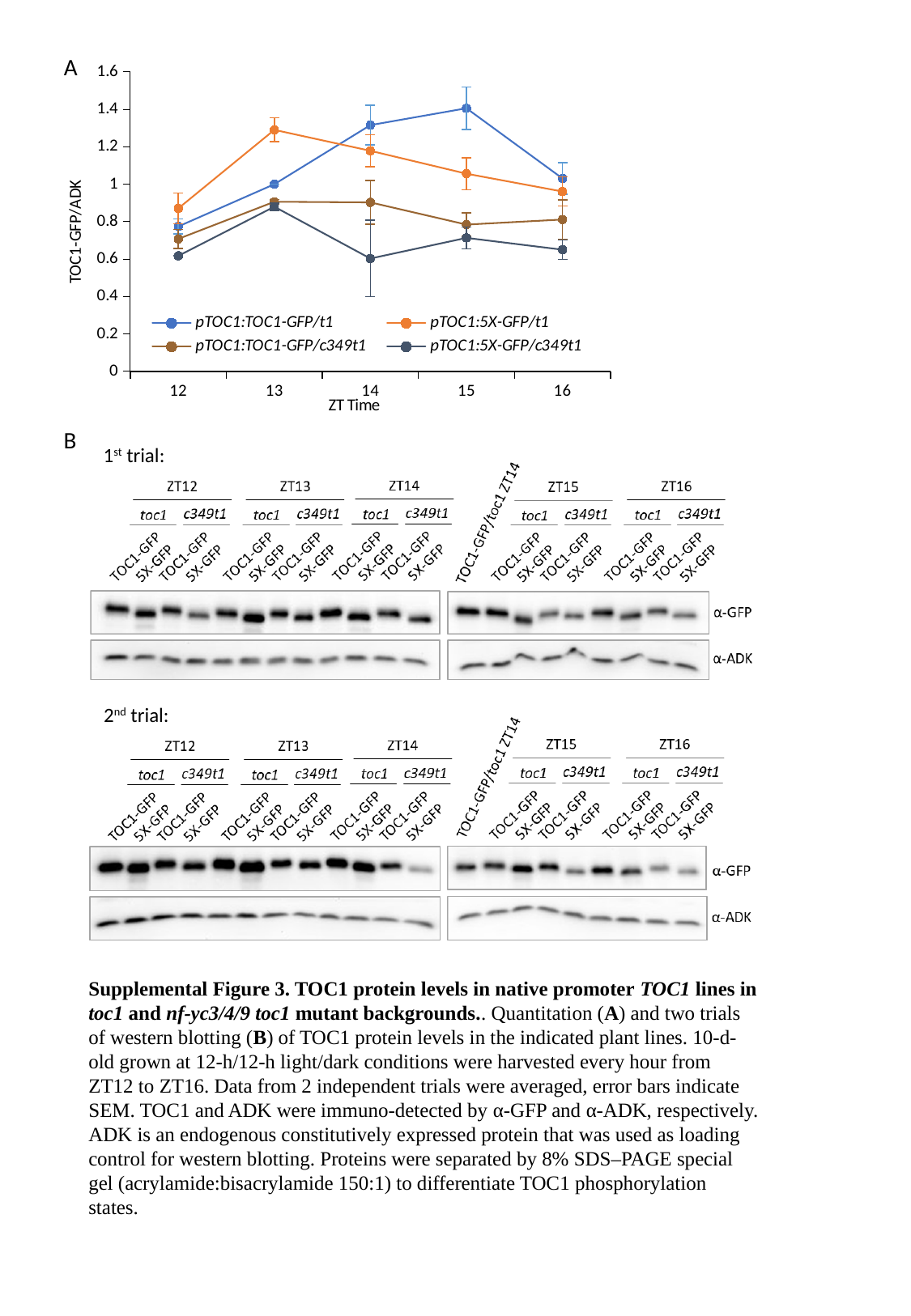

A
#### Chart
| Category | pTOC1:TOC1-GFP/t1 | pTOC1:5X-GFP/t1 | pTOC1:TOC1-GFP/c349t1 | pTOC1:5X-GFP/c349t1 |
|---|---|---|---|---|
| 12 | 0.7743397391155826 | 0.8706236528750788 | 0.7078838497094001 | 0.617255387326209 |
| 13 | 1.0 | 1.2904998430790804 | 0.9060974890426747 | 0.8796538734967054 |
| 14 | 1.3158309070233787 | 1.1783764797043859 | 0.9027711642686898 | 0.6029678109106278 |
| 15 | 1.4058956403299867 | 1.0560684179476256 | 0.78416837524294 | 0.7132572452976993 |
| 16 | 1.0307905331844616 | 0.9607483083734669 | 0.8112279789867225 | 0.65019520102462 |B
1^st^ trial:
2^nd^ trial:
Supplemental Figure 3. TOC1 protein levels in native promoter TOC1 lines in toc1 and nf-yc3/4/9 toc1 mutant backgrounds.. Quantitation (A) and two trials of western blotting (B) of TOC1 protein levels in the indicated plant lines. 10-d-old grown at 12-h/12-h light/dark conditions were harvested every hour from ZT12 to ZT16. Data from 2 independent trials were averaged, error bars indicate SEM. TOC1 and ADK were immuno-detected by α-GFP and α-ADK, respectively. ADK is an endogenous constitutively expressed protein that was used as loading control for western blotting. Proteins were separated by 8% SDS–PAGE special gel (acrylamide:bisacrylamide 150:1) to differentiate TOC1 phosphorylation states.

### Slide 4
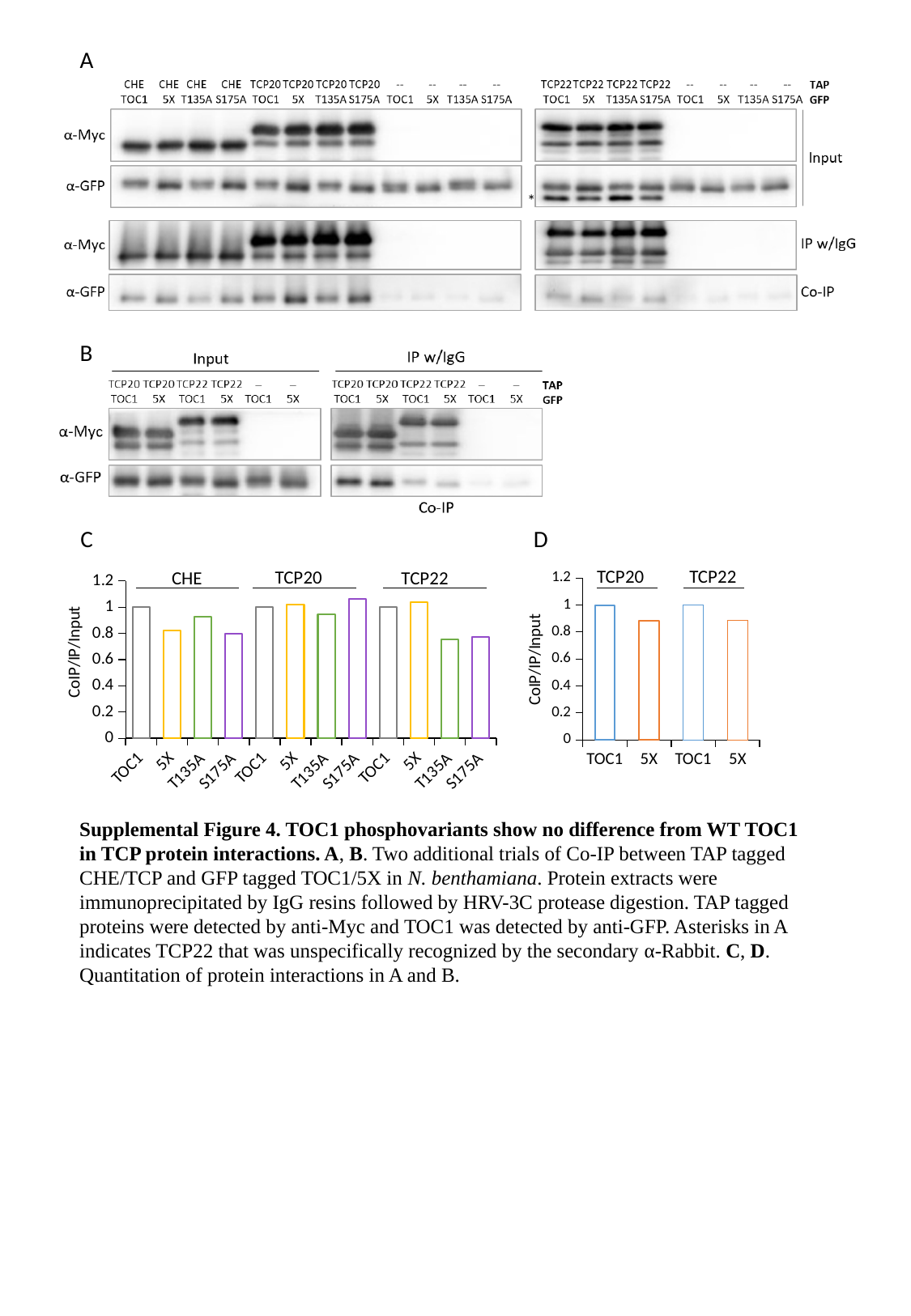

A
*
B
C
D
TCP20
TCP22
TCP20
CHE
TCP22
#### Chart
| Category | |
|---|---|
| TOC1 | 1.0 |
| 5X | 0.8223369093290139 |
| T135A | 0.9302500669686175 |
| S175A | 0.796947207625001 |
| TOC1 | 1.0 |
| 5X | 1.02125084495312 |
| T135A | 0.9447875065436481 |
| S175A | 1.0612303594910284 |
| TOC1 | 1.0 |
| 5X | 1.0406863750572373 |
| T135A | 0.7563807492670698 |
| S175A | 0.7717456294263807 |
#### Chart
| Category | |
|---|---|
| TOC1 | 1.0 |
| 5X | 0.886227625809114 |
| TOC1 | 1.0 |
| 5X | 0.8860148262265831 |Supplemental Figure 4. TOC1 phosphovariants show no difference from WT TOC1 in TCP protein interactions. A, B. Two additional trials of Co-IP between TAP tagged CHE/TCP and GFP tagged TOC1/5X in N. benthamiana. Protein extracts were immunoprecipitated by IgG resins followed by HRV-3C protease digestion. TAP tagged proteins were detected by anti-Myc and TOC1 was detected by anti-GFP. Asterisks in A indicates TCP22 that was unspecifically recognized by the secondary α-Rabbit. C, D. Quantitation of protein interactions in A and B.

### Slide 5
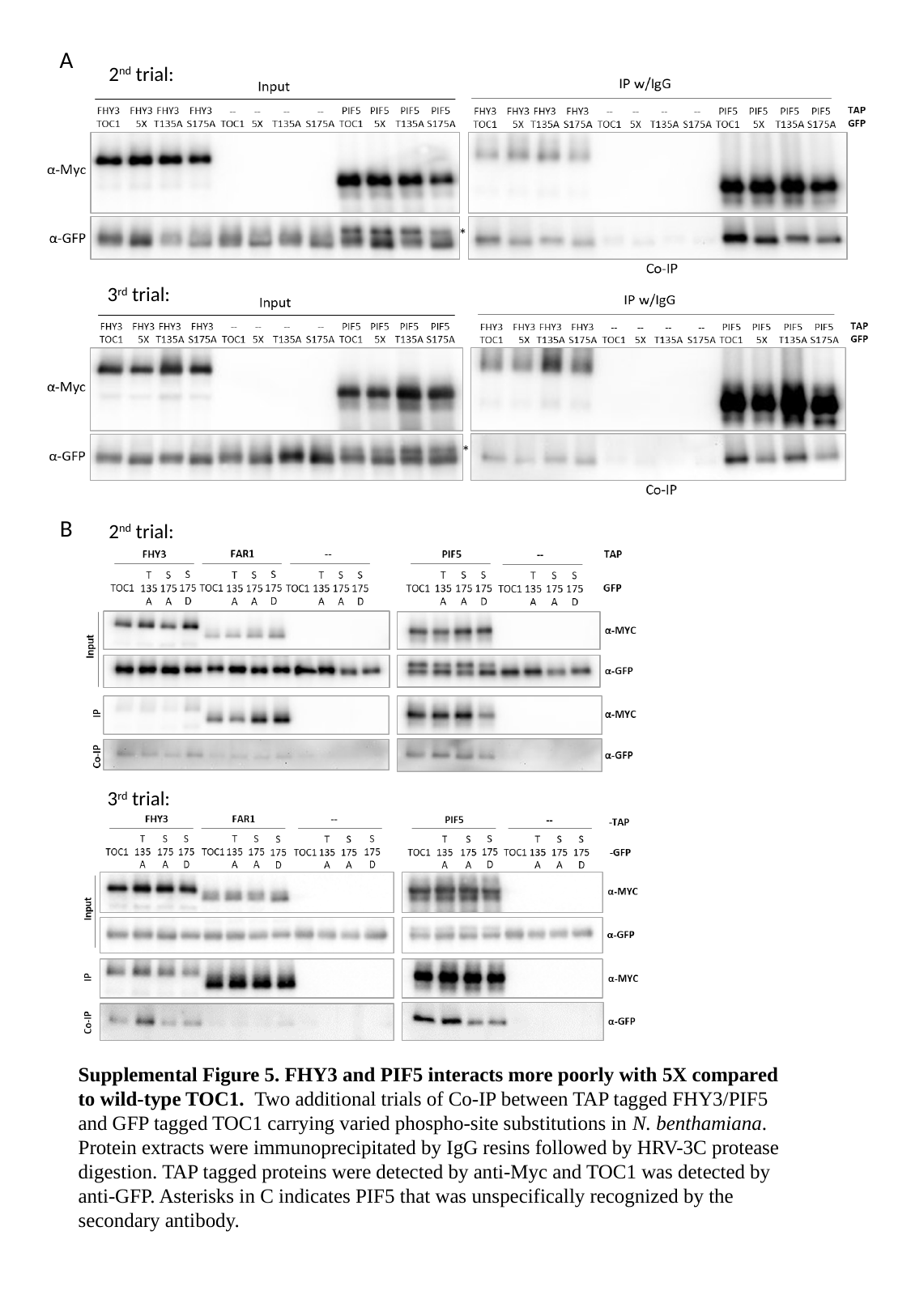

A
2^nd^ trial:
*
3^rd^ trial:
*
B
2^nd^ trial:
3^rd^ trial:
Supplemental Figure 5. FHY3 and PIF5 interacts more poorly with 5X compared to wild-type TOC1. Two additional trials of Co-IP between TAP tagged FHY3/PIF5 and GFP tagged TOC1 carrying varied phospho-site substitutions in N. benthamiana. Protein extracts were immunoprecipitated by IgG resins followed by HRV-3C protease digestion. TAP tagged proteins were detected by anti-Myc and TOC1 was detected by anti-GFP. Asterisks in C indicates PIF5 that was unspecifically recognized by the secondary antibody.

### Slide 6
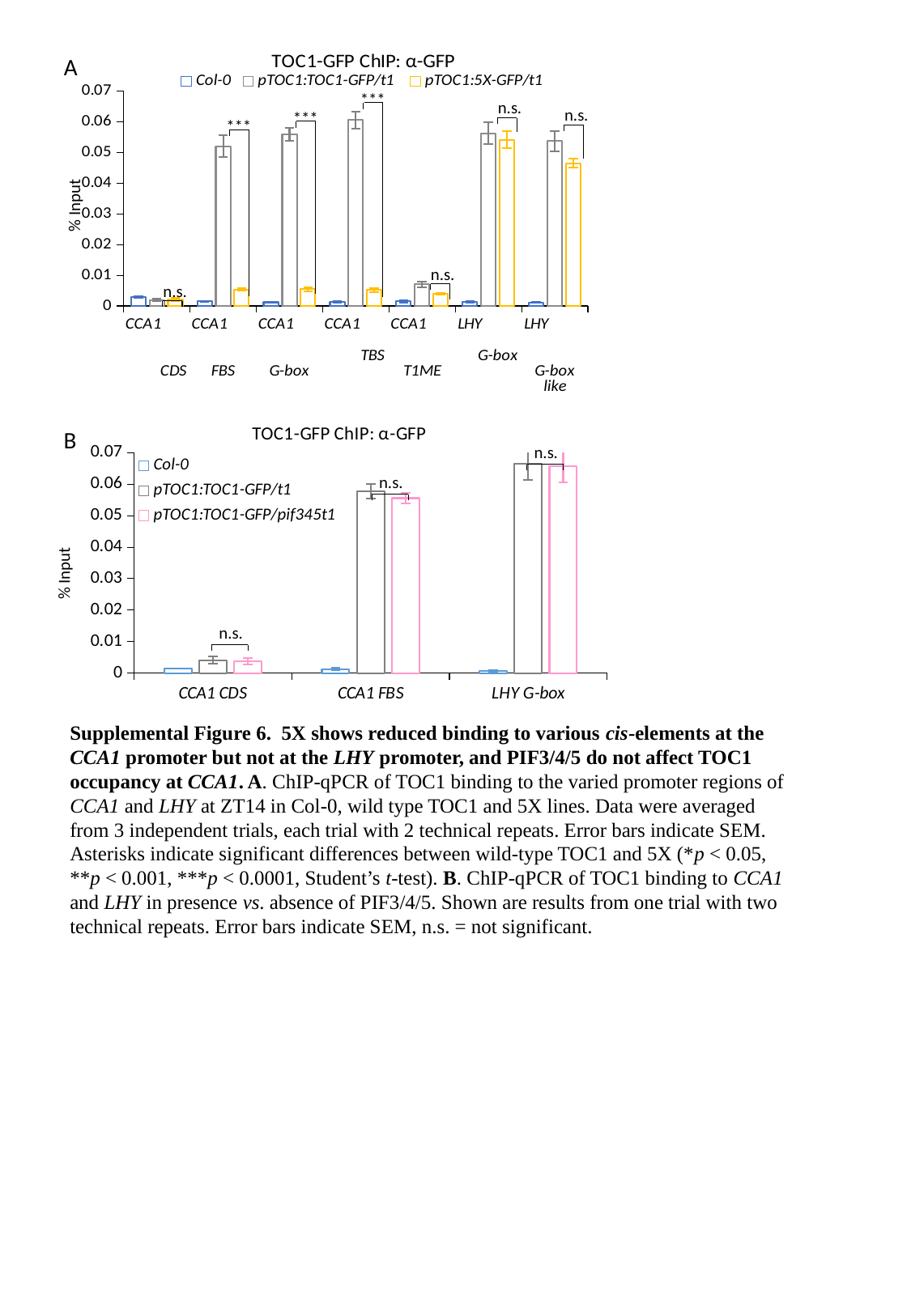

#### Chart: TOC1-GFP ChIP: α-GFP
| Category | Col-0 | pTOC1:TOC1-GFP/t1 | pTOC1:5X-GFP/t1 |
|---|---|---|---|
| CCA1 CDS | 0.0029731832664745624 | 0.0019493135015736373 | 0.0021325141028074937 |
| CCA1 FBS | 0.0014788673600884827 | 0.05209106717573645 | 0.0053644070268319835 |
| CCA1 G-box | 0.0011932608729416868 | 0.05602537001443634 | 0.005482375888466726 |
| CCA1 TBS | 0.0013494302049800767 | 0.0605489766018335 | 0.005167963035422377 |
| CCA1 T1ME | 0.001493981932249651 | 0.007032213837016177 | 0.004020751696406061 |
| LHY G-box | 0.0013400554094004109 | 0.056244950415597315 | 0.054156008469177404 |
| LHY G-box like | 0.0011359523863209884 | 0.05373148094988859 | 0.046452274563008635 |***
n.s.
n.s.
***
***
% Input
n.s.
n.s.
A
#### Chart: TOC1-GFP ChIP: α-GFP
| Category | Col-0 | pTOC1:TOC1-GFP/t1 | pTOC1:TOC1-GFP/pif345t1 |
|---|---|---|---|
| CCA1 CDS | 0.001450342437121264 | 0.004099128659365751 | 0.003702706300195111 |
| CCA1 FBS | 0.0012801052295336017 | 0.05777296578864149 | 0.055584119700084755 |
| LHY G-box | 0.0006640502809804577 | 0.06651277771302175 | 0.06572866098426704 |B
n.s.
n.s.
n.s.
Supplemental Figure 6. 5X shows reduced binding to various cis-elements at the CCA1 promoter but not at the LHY promoter, and PIF3/4/5 do not affect TOC1 occupancy at CCA1. A. ChIP-qPCR of TOC1 binding to the varied promoter regions of CCA1 and LHY at ZT14 in Col-0, wild type TOC1 and 5X lines. Data were averaged from 3 independent trials, each trial with 2 technical repeats. Error bars indicate SEM. Asterisks indicate significant differences between wild-type TOC1 and 5X (*p < 0.05, **p < 0.001, ***p < 0.0001, Student’s t-test). B. ChIP-qPCR of TOC1 binding to CCA1 and LHY in presence vs. absence of PIF3/4/5. Shown are results from one trial with two technical repeats. Error bars indicate SEM, n.s. = not significant.

### Slide 7
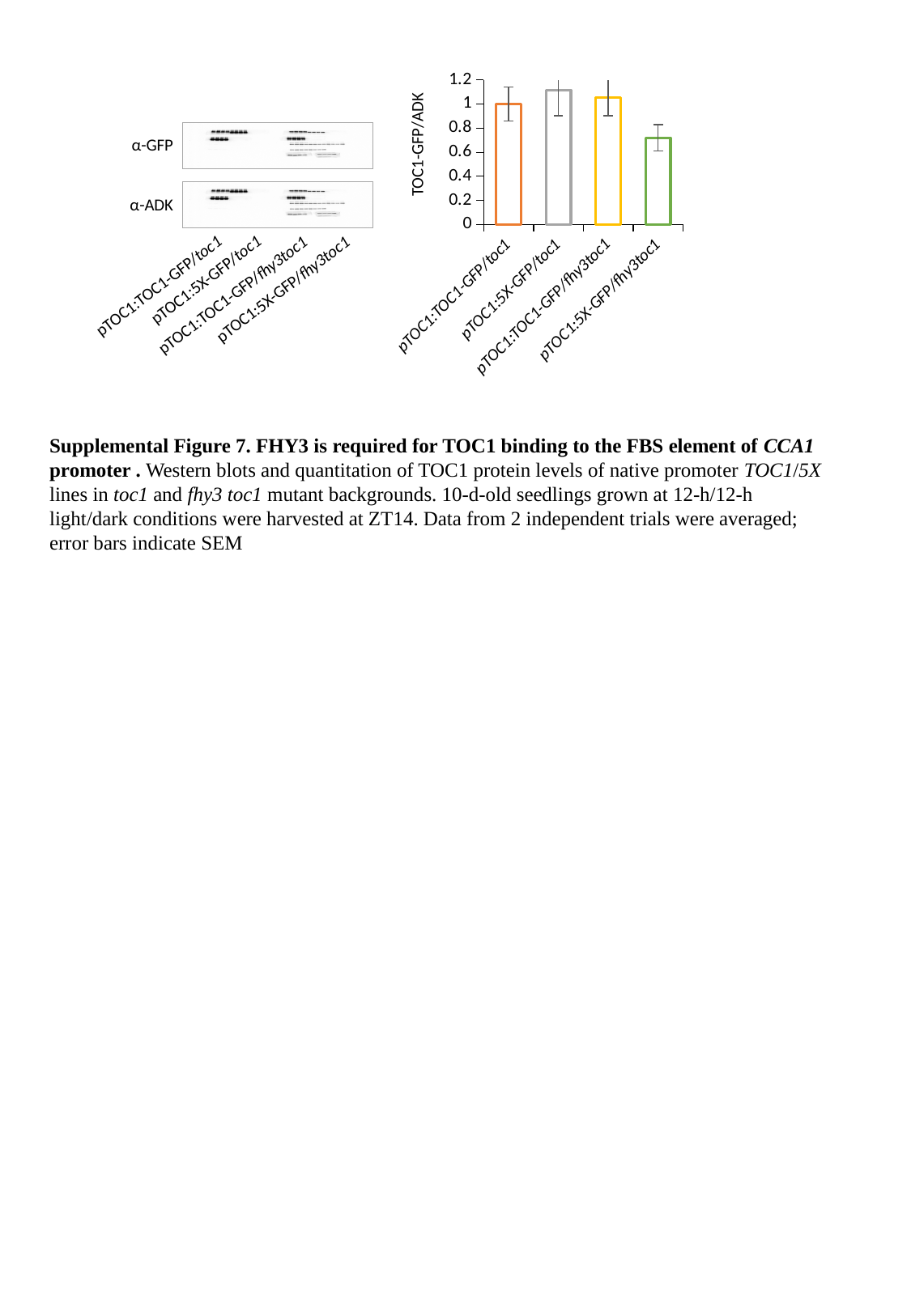

#### Chart
| Category | |
|---|---|
| pTOC1:TOC1-GFP/toc1 | 1.0 |
| pTOC1:5X-GFP/toc1 | 1.1154933527566466 |
| pTOC1:TOC1-GFP/fhy3toc1 | 1.0569273319574803 |
| pTOC1:5X-GFP/fhy3toc1 | 0.7197145648860227 |
α-GFP
α-ADK
pTOC1:5X-GFP/toc1
pTOC1:TOC1-GFP/toc1
pTOC1:5X-GFP/fhy3toc1
pTOC1:TOC1-GFP/fhy3toc1
Supplemental Figure 7. FHY3 is required for TOC1 binding to the FBS element of CCA1 promoter . Western blots and quantitation of TOC1 protein levels of native promoter TOC1/5X lines in toc1 and fhy3 toc1 mutant backgrounds. 10-d-old seedlings grown at 12-h/12-h light/dark conditions were harvested at ZT14. Data from 2 independent trials were averaged; error bars indicate SEM

### Slide 8
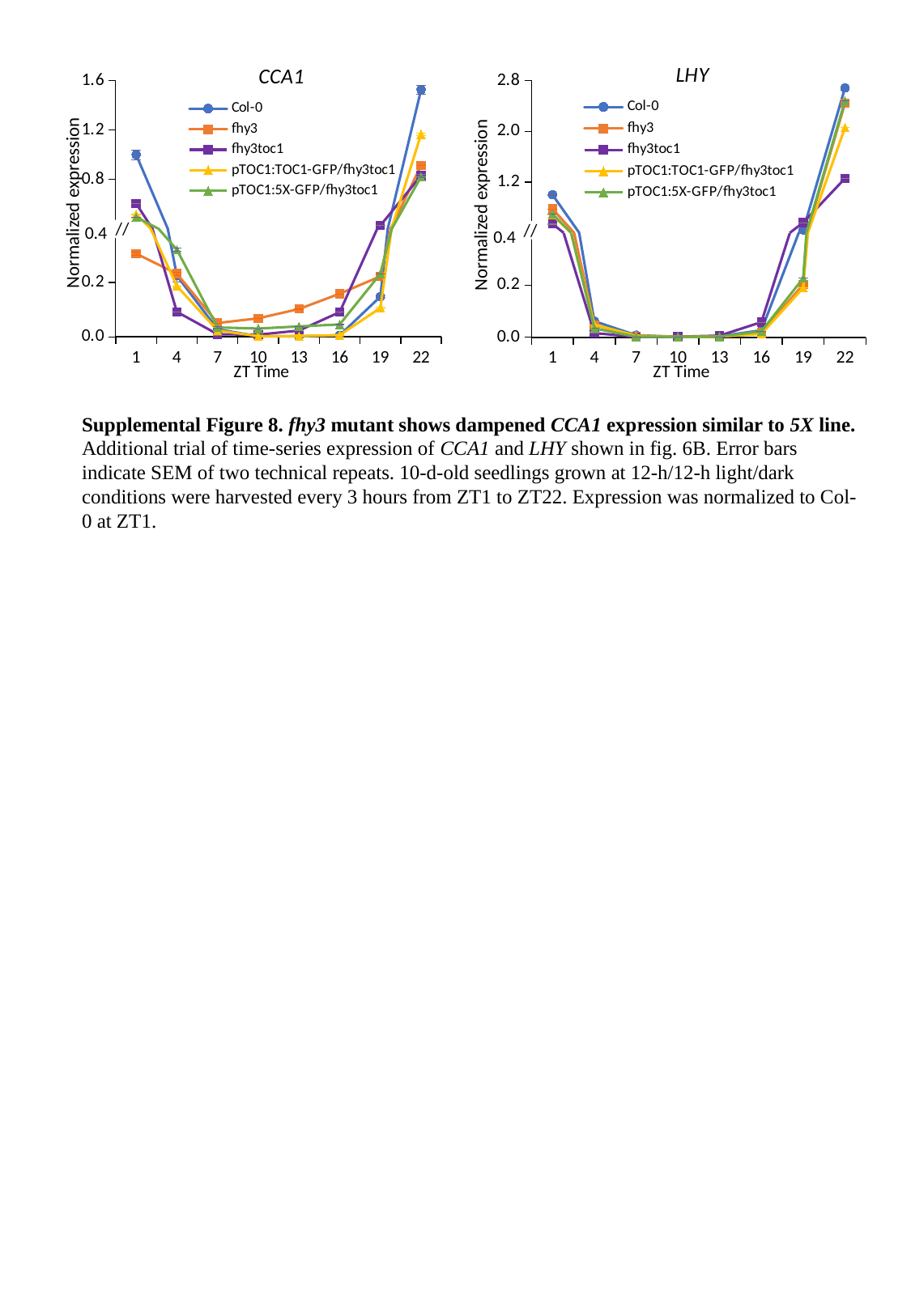

#### Chart: LHY
| Category | Col-0 | fhy3 | fhy3toc1 | pTOC1:TOC1-GFP/fhy3toc1 | pTOC1:5X-GFP/fhy3toc1 |
|---|---|---|---|---|---|
| 1 | 0.9999999999999998 | 0.776743130175046 | 0.539820140026683 | 0.6810304490369453 | 0.6958436701085842 |
| 4 | 0.06215311079911772 | 0.04824333073343519 | 0.01656887010898015 | 0.0490959763555633 | 0.034532819451431815 |
| 7 | 0.00836010910318723 | 0.0036206655799088097 | 0.0015729700089550155 | 0.006763913795635422 | 0.0024823780336563047 |
| 10 | 0.0022841138224417934 | 0.003065650137749725 | 0.0025704058523832586 | 0.002040950272268918 | 0.0025579521476239526 |
| 13 | 0.0042229968172506705 | 0.003454736559220948 | 0.006535450219823163 | 0.002832391098134374 | 0.0031520420139168753 |
| 16 | 0.027127246605896032 | 0.017834961226753068 | 0.05852798973041469 | 0.013228768279980842 | 0.02317297172028915 |
| 19 | 0.44603767025788277 | 0.20268804317629882 | 0.5615302838825424 | 0.19014965974098785 | 0.22714570384613442 |
| 22 | 2.686021964173478 | 2.44478341386713 | 1.258713575946266 | 2.061555515138253 | 2.4825416088968306 |
#### Chart: CCA1
| Category | Col-0 | fhy3 | fhy3toc1 | pTOC1:TOC1-GFP/fhy3toc1 | pTOC1:5X-GFP/fhy3toc1 |
|---|---|---|---|---|---|
| 1 | 0.9999999999999999 | 0.3059075442350935 | 0.6024133543557947 | 0.5151725873763413 | 0.4918499459107143 |
| 4 | 0.22480718342455486 | 0.23217813146218214 | 0.09198616420762881 | 0.18713941195380726 | 0.320741348645452 |
| 7 | 0.027845236546072467 | 0.05093787731375909 | 0.009576803645473642 | 0.02313223301441133 | 0.034617577497411706 |
| 10 | 0.0018292904691172283 | 0.06810940598415427 | 0.008076549119317893 | 0.002542655339725353 | 0.03040664189230078 |
| 13 | 0.0021278658610696596 | 0.10234418626433654 | 0.02314842618779327 | 0.002748138457818538 | 0.03832320805283823 |
| 16 | 0.00567527332349314 | 0.1586514291502815 | 0.09074606863517787 | 0.006295301757940606 | 0.04569096916929576 |
| 19 | 0.14796484458646683 | 0.22266759195025299 | 0.42366794590064455 | 0.10627421761068129 | 0.23169737050990022 |
| 22 | 1.527633242351897 | 0.9120079053478451 | 0.8295406698654479 | 1.1660330604426101 | 0.8202815156318021 |Normalized expression
Normalized expression
#### Chart
| Category | Col-0 | fhy3 | fhy3toc1 | pTOC1:TOC1-GFP/fhy3toc1 | pTOC1:5X-GFP/fhy3toc1 |
|---|---|---|---|---|---|
| 1 | 0.9999999999999999 | 0.3059075442350935 | 0.6024133543557947 | 0.5151725873763413 | 0.4918499459107143 |
| 4 | 0.22480718342455486 | 0.23217813146218214 | 0.09198616420762881 | 0.18713941195380726 | 0.320741348645452 |
| 7 | 0.027845236546072467 | 0.05093787731375909 | 0.009576803645473642 | 0.02313223301441133 | 0.034617577497411706 |
| 10 | 0.0018292904691172283 | 0.06810940598415427 | 0.008076549119317893 | 0.002542655339725353 | 0.03040664189230078 |
| 13 | 0.0021278658610696596 | 0.10234418626433654 | 0.02314842618779327 | 0.002748138457818538 | 0.03832320805283823 |
| 16 | 0.00567527332349314 | 0.1586514291502815 | 0.09074606863517787 | 0.006295301757940606 | 0.04569096916929576 |
| 19 | 0.14796484458646683 | 0.22266759195025299 | 0.42366794590064455 | 0.10627421761068129 | 0.23169737050990022 |
| 22 | 1.527633242351897 | 0.9120079053478451 | 0.8295406698654479 | 1.1660330604426101 | 0.8202815156318021 |
#### Chart
| Category | Col-0 | fhy3 | fhy3toc1 | pTOC1:TOC1-GFP/fhy3toc1 | pTOC1:5X-GFP/fhy3toc1 |
|---|---|---|---|---|---|
| 1 | 0.9999999999999998 | 0.776743130175046 | 0.539820140026683 | 0.6810304490369453 | 0.6958436701085842 |
| 4 | 0.06215311079911772 | 0.04824333073343519 | 0.01656887010898015 | 0.0490959763555633 | 0.034532819451431815 |
| 7 | 0.00836010910318723 | 0.0036206655799088097 | 0.0015729700089550155 | 0.006763913795635422 | 0.0024823780336563047 |
| 10 | 0.0022841138224417934 | 0.003065650137749725 | 0.0025704058523832586 | 0.002040950272268918 | 0.0025579521476239526 |
| 13 | 0.0042229968172506705 | 0.003454736559220948 | 0.006535450219823163 | 0.002832391098134374 | 0.0031520420139168753 |
| 16 | 0.027127246605896032 | 0.017834961226753068 | 0.05852798973041469 | 0.013228768279980842 | 0.02317297172028915 |
| 19 | 0.44603767025788277 | 0.20268804317629882 | 0.5615302838825424 | 0.19014965974098785 | 0.22714570384613442 |
| 22 | 2.686021964173478 | 2.44478341386713 | 1.258713575946266 | 2.061555515138253 | 2.4825416088968306 |//
//
0.4
0.4
Supplemental Figure 8. fhy3 mutant shows dampened CCA1 expression similar to 5X line. Additional trial of time-series expression of CCA1 and LHY shown in fig. 6B. Error bars indicate SEM of two technical repeats. 10-d-old seedlings grown at 12-h/12-h light/dark conditions were harvested every 3 hours from ZT1 to ZT22. Expression was normalized to Col-0 at ZT1.

### Slide 9
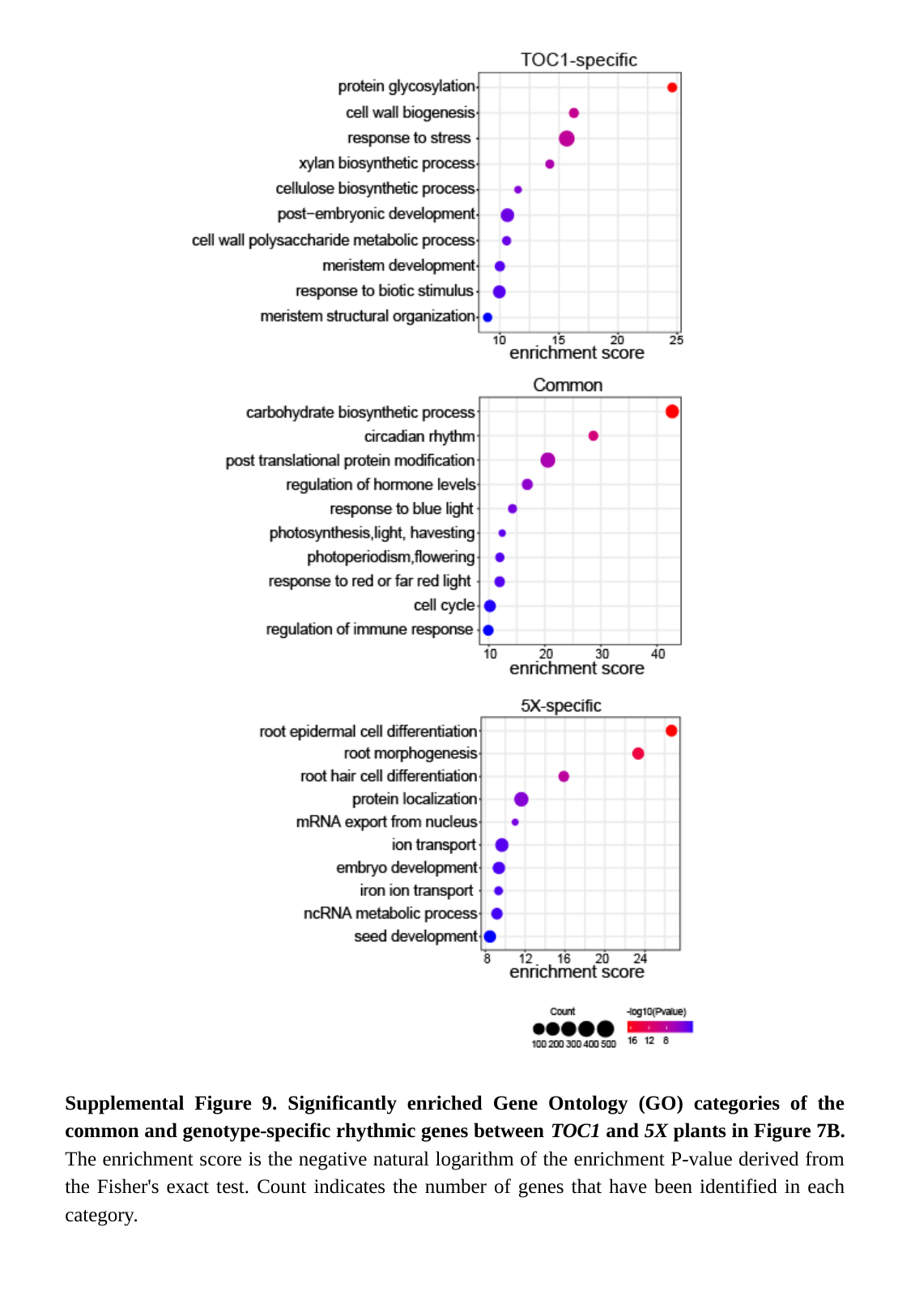

Supplemental Figure 9. Significantly enriched Gene Ontology (GO) categories of the common and genotype-specific rhythmic genes between TOC1 and 5X plants in Figure 7B. The enrichment score is the negative natural logarithm of the enrichment P-value derived from the Fisher's exact test. Count indicates the number of genes that have been identified in each category.

### Slide 10
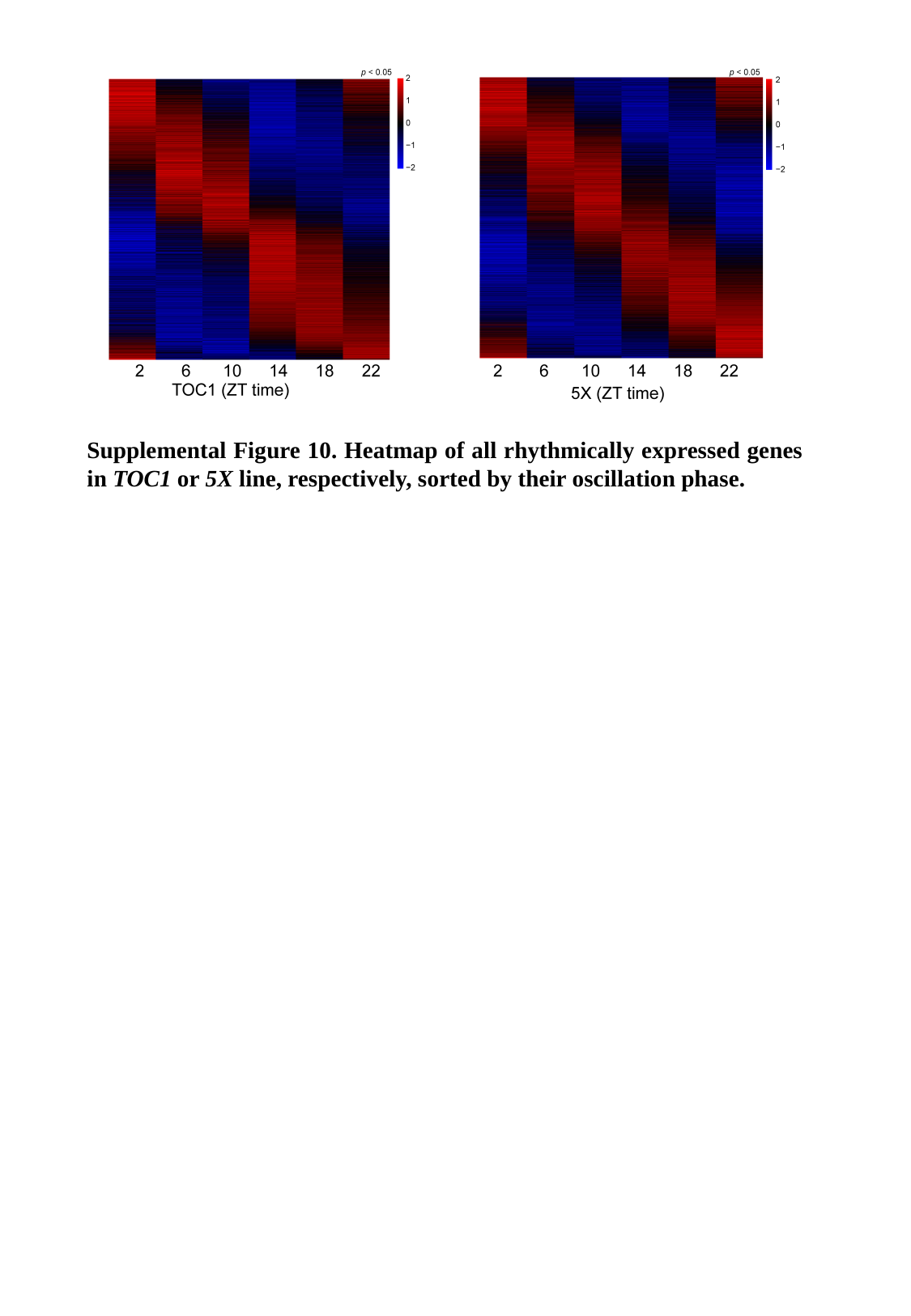

Supplemental Figure 10. Heatmap of all rhythmically expressed genes in TOC1 or 5X line, respectively, sorted by their oscillation phase.

### Slide 11
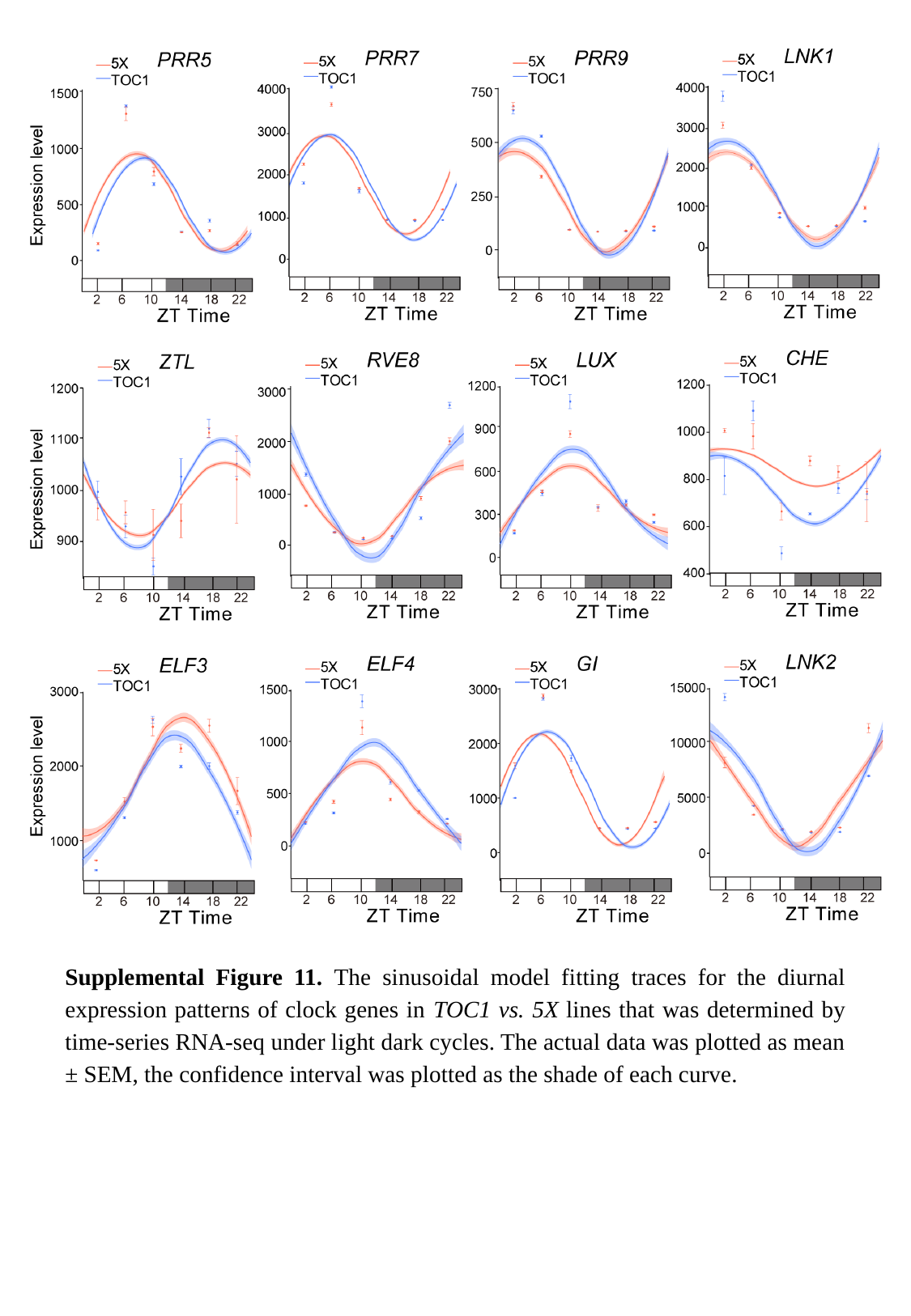

Supplemental Figure 11. The sinusoidal model fitting traces for the diurnal expression patterns of clock genes in TOC1 vs. 5X lines that was determined by time-series RNA-seq under light dark cycles. The actual data was plotted as mean ± SEM, the confidence interval was plotted as the shade of each curve.

### Slide 12
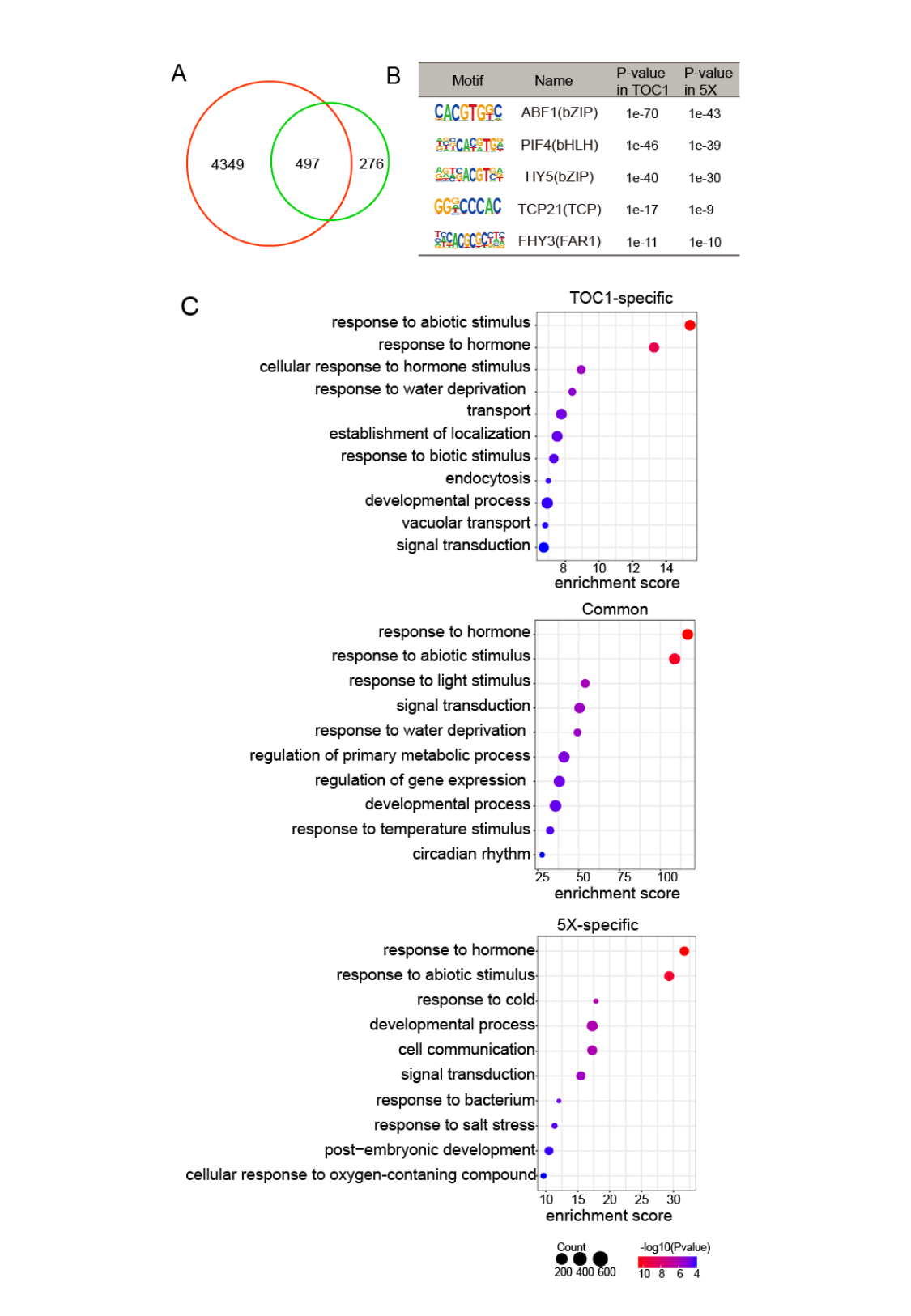

### Slide 13
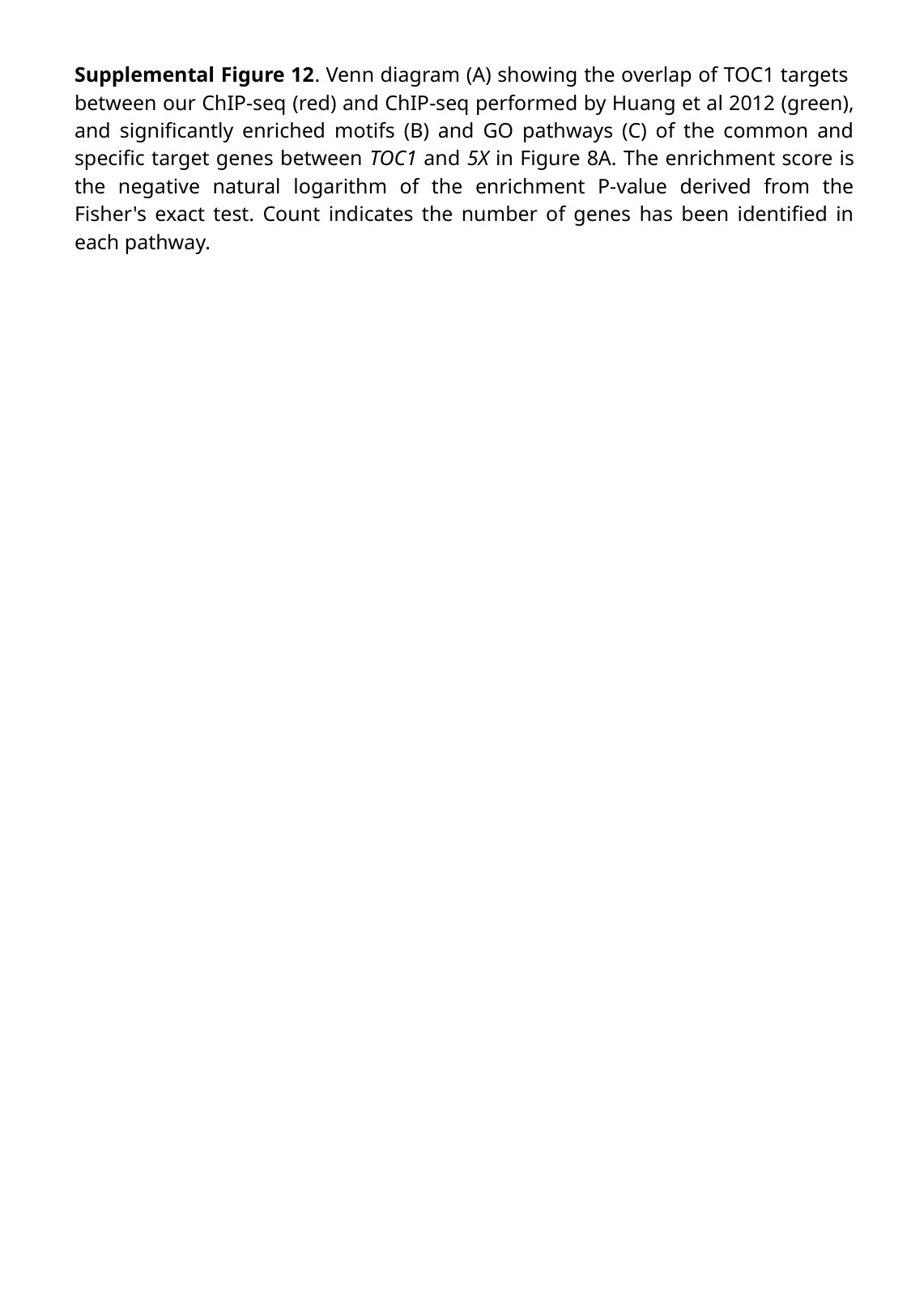

Supplemental Figure 12. Venn diagram (A) showing the overlap of TOC1 targets between our ChIP-seq (red) and ChIP-seq performed by Huang et al 2012 (green), and significantly enriched motifs (B) and GO pathways (C) of the common and specific target genes between TOC1 and 5X in Figure 8A. The enrichment score is the negative natural logarithm of the enrichment P-value derived from the Fisher's exact test. Count indicates the number of genes has been identified in each pathway.

### Slide 14
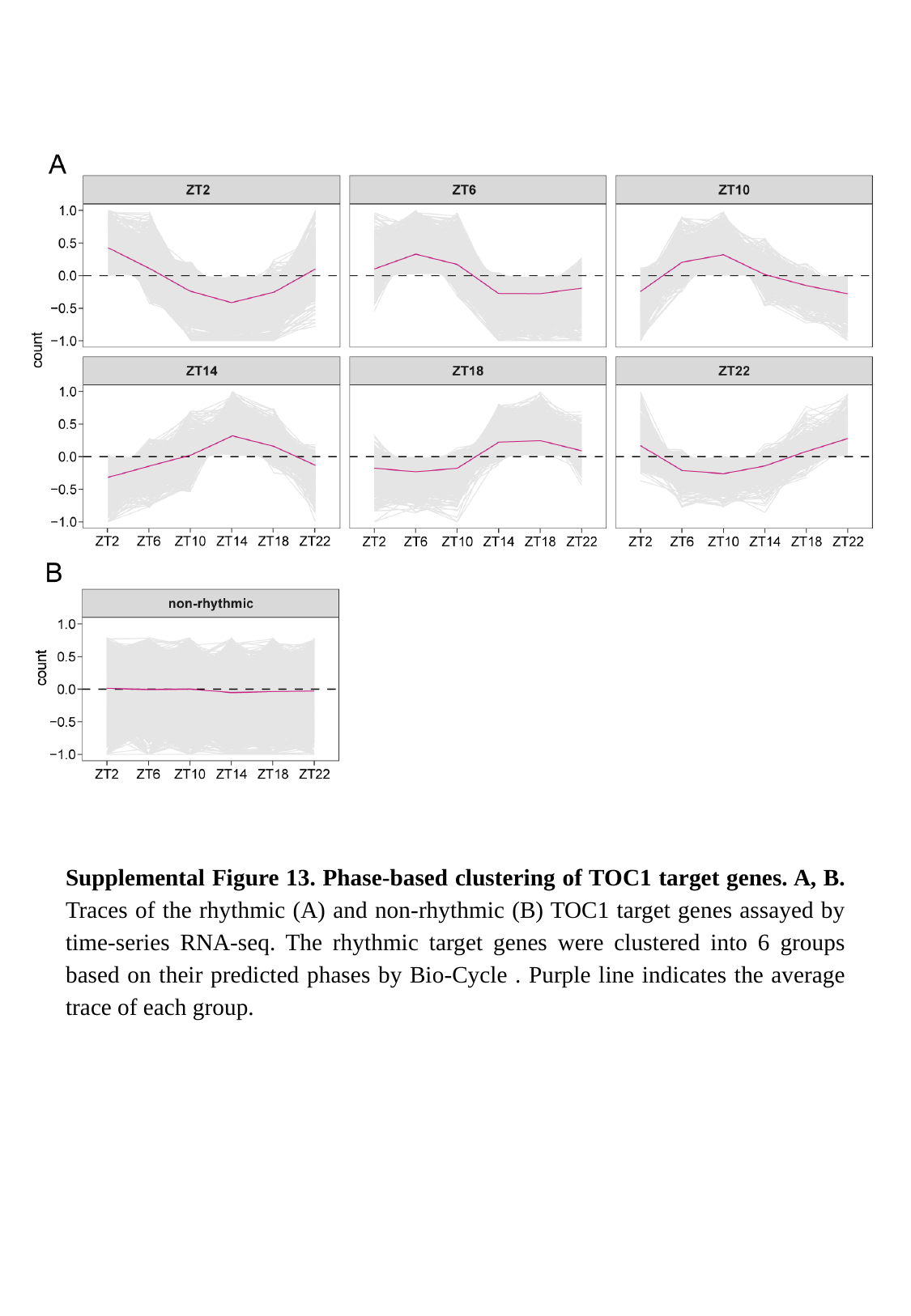

Supplemental Figure 13. Phase-based clustering of TOC1 target genes. A, B. Traces of the rhythmic (A) and non-rhythmic (B) TOC1 target genes assayed by time-series RNA-seq. The rhythmic target genes were clustered into 6 groups based on their predicted phases by Bio-Cycle . Purple line indicates the average trace of each group.

### Slide 15
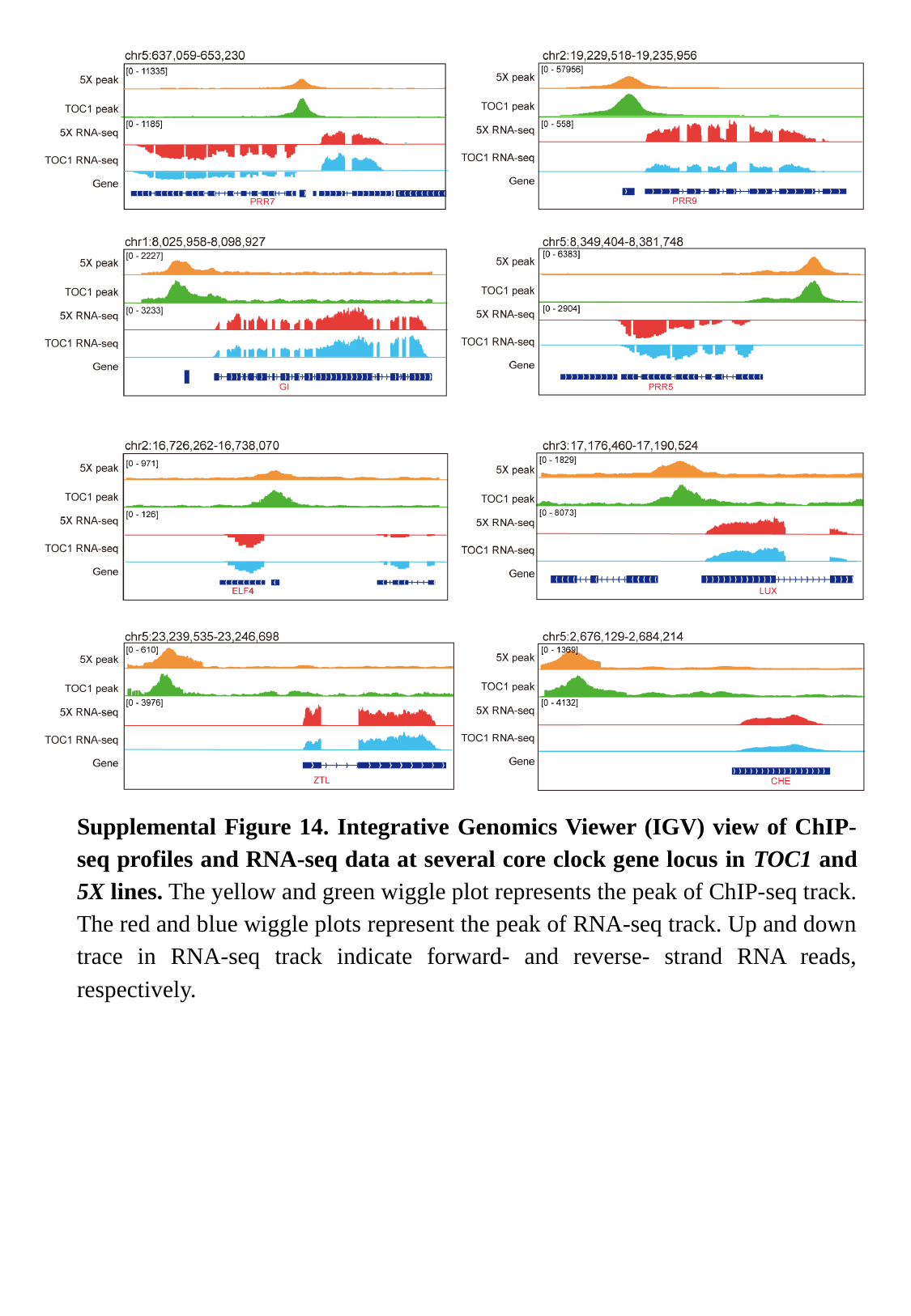

Supplemental Figure 14. Integrative Genomics Viewer (IGV) view of ChIP-seq profiles and RNA-seq data at several core clock gene locus in TOC1 and 5X lines. The yellow and green wiggle plot represents the peak of ChIP-seq track. The red and blue wiggle plots represent the peak of RNA-seq track. Up and down trace in RNA-seq track indicate forward- and reverse- strand RNA reads, respectively.
